# Tonic pain revalues associative memories of phasic pain

**DOI:** 10.1101/2025.03.04.641525

**Authors:** Danielle Hewitt, Shuangyi Tong, Sarah Schreiber, Ben Seymour

## Abstract

Tonic pain is proposed to adapt protective behaviours during recovery from injury. A key untested prediction of this homeostatic model is that it appropriately reshapes internal representations of phasic pain. We investigated whether lateralised tonic pain modulates phasic pain-predictive cues on that side. Using a virtual-reality Pavlovian revaluation paradigm, we assessed physiological and neural conditioned responses with EEG in theta, alpha, and beta frequency bands. Pain-predictive cues elicited neural enhanced alpha and beta suppression and increased pupil diameter during conditioning acquisition. Critically, tonic pain revalued phasic conditioned responses during extinction, with reduced midfrontal theta synchronisation when the laterality of tonic pain was congruent with predicted phasic pain. Greater tonic pain unpleasantness also enhanced posterior beta suppression for congruent cues. These findings provide evidence for an internal representation of cue-pain associations that is topographically modulated by tonic pain, suggesting that tonic pain actively reconfigures pain predictions, enabling anticipatory protective behaviours.

## 1. Introduction

Classical notions of pain propose that tonic and phasic pain subserve distinct functions in adaptive behaviour. The role of phasic pain in rapid harm detection and nocifensive behaviour is well characterised, and greatly amplified by the capacity for learning, allowing anticipatory responding and avoidance (Pavlovian and instrumental conditioning) ^1^. In contrast, tonic pain is thought to enhance hyper-protective and recuperative behaviour appropriate to the recovery from injury, given the state of vulnerability and changed homeostatic demands ^2,3^. This has led to the concept of tonic pain as a homeostatic state, as it relates to the widespread control of physiology and behaviour prioritising the maintenance of bodily integrity. A cardinal (but untested) prediction of this homeostatic model of tonic pain function is that it reshapes internal representations (memories) of phasic pain.

This relationship is well established in the reward literature, where homeostatic states such as hunger or thirst reshape associative memories of food or drinks. In the classical ‘devaluation’ paradigm, animals are trained to associate a novel cue with a particular food whilst hungry, but (Pavlovian) responding for that food is diminished when sated after feeding ^4,5^. Reshaping specific to the reward type is taken as evidence of an internal representation of the reward — so-called ‘model-based’ learning — which is clearly adaptive as motivation is appropriately modulated by homeostatic energy requirements ^6^. Applying this logic to pain would be similarly adaptive; for instance, a painful injury to the right arm should amplify previously learned associative memories for things that might cause phasic pain to the area, as an injured region will be more susceptible to further damage. Here, we aim to test the homeostatic revaluation hypothesis by showing that lateralised tonic pain specifically modulates associative responses to a phasic pain stimulus on that side.

Demonstrating the reshaping of internal representations of pain faces two methodological challenges. The first is imaging these representations. This can be addressed using electroencephalography (EEG), which can reveal distinct neural signatures shaped by predictions and experiences. Aversive conditioned stimuli (sounds or electric shocks) modulate EEG cortical oscillations in alpha (8–12 Hz) and theta (4–7 Hz) bands ^7–9^, and pain memories specifically elicit suppressed alpha activity over parietal regions, implying somatomotor representations appropriate to pain-predictive cues ^10,11^. The second challenge is embedding the associative memory into a sufficiently threatening context, to indue the behavioural relevance of the memory when related to the tonic pain. This can be solved using virtual reality (VR), where a conditioning task can be set within an ecologically realistic and immersive context.

We therefore designed a classical Pavlovian revaluation paradigm, using lateralised phasic pain conditioning and externally applied tonic pain as a proxy for a homeostatic manipulation, implemented in an EEG-VR design. During the task, we recorded changes in EEG conditioned responses in theta, alpha and beta (16–24 Hz) frequency bands alongside physiological arousal indicators (pupil diameter and fixations). In accordance with our predictions, we found heightened neural conditioned responses to pain-predictive compared to neutral cues during conditioning acquisition, indexed by enhanced alpha and beta suppression and increased pupil diameter. In support of our primary revaluation hypothesis, we found that that lateralised tonic pain revalued phasic neural conditioned responses related to that side.

## 2. Results

Twenty-six human volunteers participated fully in the task. In the first stage, participants underwent Pavlovian trace conditioning to left- or right-sided painful shocks. The conditioned stimuli were animated foxes with different coloured hats within a VR game, with one colour designated the CS+ for right-sided pain, another colour for left-sided pain, and a third colour as a neutral cue (see Figure 1). The foxes appeared within the VR game for 0.8 s, followed by a 2 s waiting period where no fox was present, after which a rock was ‘thrown’ towards the subject slightly to the left or right side (followed by pain to the appropriate side), or in the case of the neutral cue, a central moving rock that fell short. The unconditioned stimulus was therefore the compound of incoming rock and pain. In the second stage, tonic pain was applied to one side via an upper arm pressure cuff. The cues were then shown in extinction (such that the subject never simultaneously experienced phasic and tonic pain – a core requirement for demonstrating de-/re-valuation).

**Figure 1.**
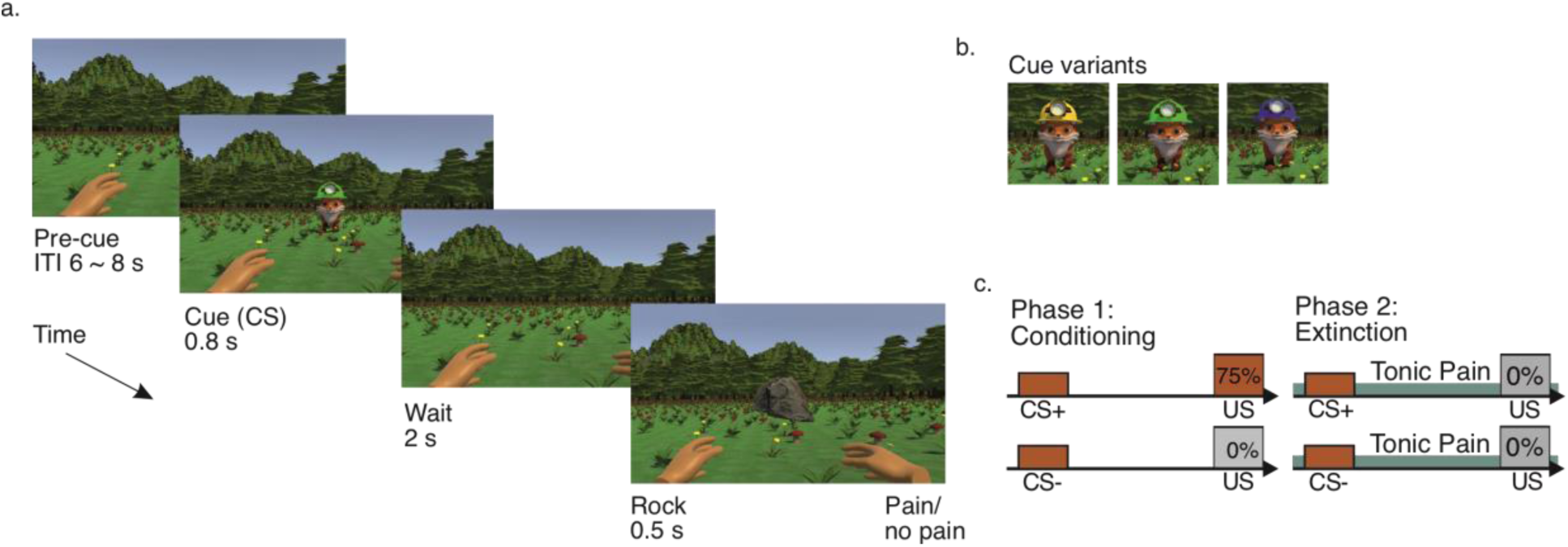
Pavlovian revaluation paradigm schematic. a. At the start of each trial, an animated fox wearing a green, yellow, or blue hat (conditioned stimulus, CS; each colour corresponding to a cue condition) appeared in the centre of the visual field for 0.8 s, followed by a 2 s waiting period. Subsequently, a rock appeared and travelled with a left, right, or central trajectory towards the participant’s hands (left/right) or the ground (middle), with a looming duration of 0.5 s, accompanied by a binaural sound that increased in volume as the object approached. Trials were followed by a random intertrial interval of 6-8 s. b. Looming direction was determined by the fox’s hat colour, with cues corresponding to each trial type counterbalanced for each participant. The unconditioned stimulus was therefore the compound of incoming rock and pain. c. Pavlovian (Trace) conditioning schematic. In phase 1 conditioning blocks, pain-related CS+ cues were followed by a 75% chance of experiencing phasic pain. Neutral CS- cues were followed by a 0% chance of experiencing pain. In phase 2 extinction blocks, no cues were followed by phasic pain. Tonic pain was delivered throughout the extinction block using an upper arm pressure cuff.

In basic analyses, there were no effects of laterality on pain ratings. Phasic electrical and tonic pressure stimuli were rated as painful (phasic 4.45±1.28, tonic 3.61±1.09) and unpleasant (phasic 4.62±1.36, tonic 4.35±1.31) throughout the experiment. As expected, there were no significant effects of laterality (left vs. right) on tonic or phasic pain (*p >* .05). The effect of time (within- and between-blocks) on pain ratings and unpleasantness was significant for both tonic and phasic pain, with reduced ratings over time and between blocks (*p <* .05; Supplementary Materials).

### 2.1 Behavioural evidence of conditioning

We found physiological evidence of conditioning of the painful cue (CS+) to the shock (US). Normalised pupil diameter over the entire trial duration showed a negative deflection after cue presentation, peaking at around 1.1s, followed by a positive deflection indicating pupil dilation until the end of the trial (Figure *2*a). Visual inspection suggested greater pupil dilation for pain-related vs. neutral trials and conditioning vs. extinction blocks (with tonic pain delivered during extinction blocks), starting after 1.8 s (during the anticipation interval), in line with literature on the temporal dynamics of conditioning and anticipation effects ^9,10^. Accordingly, the anticipation interval was divided into two 1 s segments for further analyses: early (800–1799 ms) and late (1800–2799 ms) anticipation.

**Figure 2.**
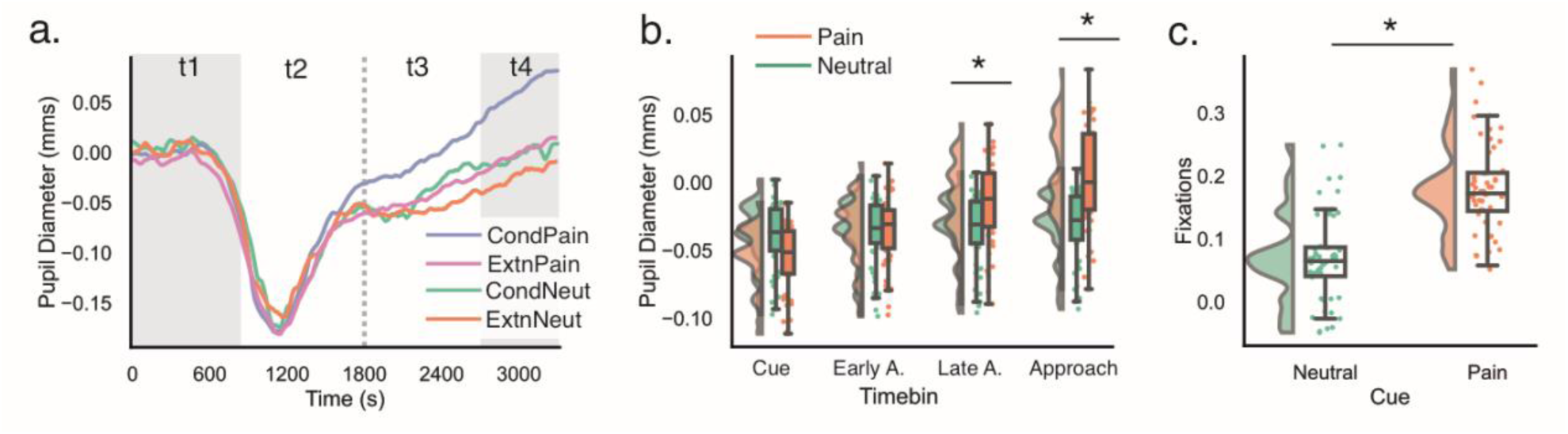
Mean pupil diameter during the trial and fixations on the approaching rock. a. Grand-average changes in pupil diameter compared to the pre-trial baseline are plotted continuously over the entire trial, with shaded areas representing CS presentation (t1) and rock approach (t4). The dotted line represents separation of the CS-US interval into early (t2) and late anticipation (t3) timebins. b. Mean pupil diameter during each of the four timebins was averaged within discrete intervals for analysis. c. Fixations during the approach period (t4) were averaged and compared for neutral and pain-related cues. Asterisks indicate significant results at p < .05 following FWE-correction.

Physiological data were entered into individual linear mixed-effects models (LMM) to assess changes as a function of block (conditioning and extinction), cue (neutral, pain-related), and measurement interval (cue, early anticipation, late anticipation, and rock approach timebins). Full results are presented in **Table 1**.

**Table 1.**
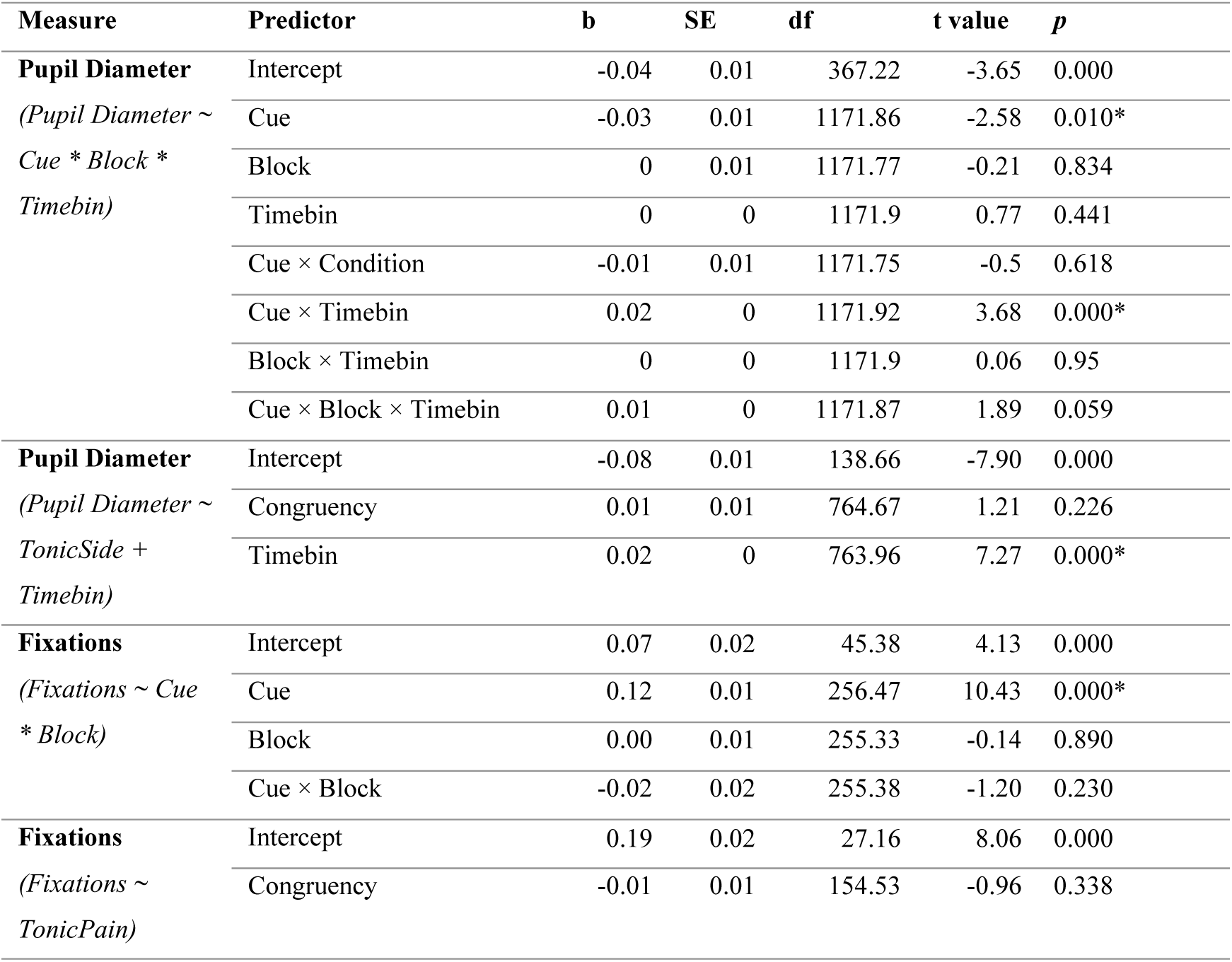
Fixed effects for physiological comparisons of interest. Asterisks denote effects that survived FWE correction (*p* < .05). Sum contrast coding was used for Block. Default (treatment) contrast coding was used for Cue (reference: neutral), measurement timebin (reference: cue) and tonic pain side (reference: incongruent).

| Measure | Predictor | b | SE | df | t value | p |
| --- | --- | --- | --- | --- | --- | --- |
| **Pupil Diameter**<br>(*Pupil Diameter ~ Cue \* Block \* Timebin*) | Intercept | -0.04 | 0.01 | 367.22 | -3.65 | 0.000 |
|  | Cue | -0.03 | 0.01 | 1171.86 | -2.58 | 0.010* |
|  | Block | 0 | 0.01 | 1171.77 | -0.21 | 0.834 |
|  | Timebin | 0 | 0 | 1171.9 | 0.77 | 0.441 |
|  | Cue × Condition | -0.01 | 0.01 | 1171.75 | -0.5 | 0.618 |
|  | Cue × Timebin | 0.02 | 0 | 1171.92 | 3.68 | 0.000* |
|  | Block × Timebin | 0 | 0 | 1171.9 | 0.06 | 0.95 |
|  | Cue × Block × Timebin | 0.01 | 0 | 1171.87 | 1.89 | 0.059 |
| **Pupil Diameter**<br>(*Pupil Diameter ~ TonicSide + Timebin*) | Intercept | -0.08 | 0.01 | 138.66 | -7.90 | 0.000 |
|  | Congruency | 0.01 | 0.01 | 764.67 | 1.21 | 0.226 |
|  | Timebin | 0.02 | 0 | 763.96 | 7.27 | 0.000* |
| **Fixations**<br>(*Fixations ~ Cue \* Block*) | Intercept | 0.07 | 0.02 | 45.38 | 4.13 | 0.000 |
|  | Cue | 0.12 | 0.01 | 256.47 | 10.43 | 0.000* |
|  | Block | 0.00 | 0.01 | 255.33 | -0.14 | 0.890 |
|  | Cue × Block | -0.02 | 0.02 | 255.38 | -1.20 | 0.230 |
| **Fixations**<br>(*Fixations ~ TonicPain*) | Intercept | 0.19 | 0.02 | 27.16 | 8.06 | 0.000 |
|  | Congruency | -0.01 | 0.01 | 154.53 | -0.96 | 0.338 |

For pupil diameter, LMMs showed a significant main effect of cue. This effect was due to more dilated pupils for pain versus neutral cues, although the corrected pairwise comparison did not reach significance (t(1180) = -1.90, *p* = .058). This result was qualified by a cue × timebin interaction. Breaking the effect down by cue, we observed greater pupil dilation for pain compared to neutral cues in late anticipation (t(1180) = -3.23, *p* = .001) and approach timebins (t(1180) = -4.07, *p* < .001; *Figure 2*b), with no difference based on cue valence during cue presentation and early anticipation (*p >* .05). The main effect of block was not significant (*p >* .05).

Fixations on the approaching rock were compared between cue types and conditions. Rock fixations accounted for 0-64% of total fixations during the approach interval. There was a statistically significant effect of cue, due to more fixations on looming objects following pain-related versus neutral cues (t(260) = -13.43, *p* < .001; *Figure 2*c). No significant effect of block was observed.

### 2.2 Neural conditioned responses

There was EEG evidence of conditioning in alpha and beta frequency bands. LMMs were fit for each frequency band and electrode cluster to assess changes as a function of block (conditioning and extinction), cue (neutral CS-, pain-related CS+), and measurement interval (cue, early anticipation and late anticipation). Full results are presented in **Table 2**.

**Table 2.** Fixed effects for EEG comparisons of interest. Asterisks denote effects that survived FWE correction (*p* < .05). Default (treatment) contrast coding was used for variables of Block (reference: conditioning), Cue (reference: neutral), and measurement timebin (reference: cue).

| Cluster | Predictor | Band |  |  |  |  |  |  |  |  |  |  |  |  |  |  |
| --- | --- | --- | --- | --- | --- | --- | --- | --- | --- | --- | --- | --- | --- | --- | --- | --- |
|  |  | Alpha |  |  |  |  | Beta |  |  |  |  | Theta |  |  |  |  |
|  |  | *b* | *SE* | *df* | *t value* | *p* | *b* | *SE* | *df* | *t value* | *p* | *b* | *SE* | *df* | *t value* | *p* |
| Frontal | Intercept | 9.79 | 2.63 | 215.08 | 3.72 | 0 | 2.91 | 1.56 | 270.21 | 1.87 | 0.063 | 27.11 | 2.8 | 442.17 | 9.69 | 0 |
|  | Cue | 2.78 | 2.65 | 2696.52 | 1.05 | 0.293 | -1.1 | 1.61 | 2756 | -0.68 | 0.496 | -2.34 | 3.05 | 2646.57 | -0.76 | 0.444 |
|  | Block | -6.16 | 3.05 | 2696.48 | -2.02 | 0.044 | -1.58 | 1.86 | 2756 | -0.85 | 0.398 | -11 | 3.51 | 2646.3 | -3.14 | **0.002\*** |
|  | Timebin | -2.12 | 1 | 2696.51 | -2.12 | 0.034 | -0.08 | 0.61 | 2756 | -0.13 | 0.899 | -8.68 | 1.15 | 2646.07 | -7.57 | **0.000\*** |
|  | Cue × Block | 2.74 | 3.73 | 2696.46 | 0.73 | 0.463 | 3.02 | 2.28 | 2756 | 1.32 | 0.186 | 4.3 | 4.31 | 2646.56 | 1 | 0.318 |
|  | Cue × Timebin | -2.65 | 1.22 | 2696.56 | -2.17 | 0.03 | -0.71 | 0.75 | 2756 | -0.95 | 0.343 | -0.31 | 1.41 | 2646.09 | -0.22 | 0.828 |
|  | Block × Timebin | 1.68 | 1.41 | 2696.64 | 1.19 | 0.234 | 0.05 | 0.86 | 2756 | 0.06 | 0.951 | 3.24 | 1.62 | 2646.23 | 2 | 0.045 |
|  | Cue × Block × Timebin | 0.84 | 1.73 | 2696.62 | 0.49 | 0.625 | -0.55 | 1.06 | 2756 | -0.52 | 0.601 | -0.76 | 1.99 | 2646.38 | -0.38 | 0.703 |
| Central | Intercept | 1.79 | 2.16 | 86.55 | 0.83 | 0.408 | -0.17 | 1.3 | 210.01 | -0.13 | 0.898 | 18.87 | 1.65 | 307.44 | 11.41 | 0 |
|  | Cue | 4.97 | 1.8 | 6410.88 | 2.76 | **0.006\*** | 2.67 | 1.3 | 6490.02 | 2.06 | 0.039 | 1.45 | 1.73 | 6393.64 | 0.84 | 0.404 |
|  | Block | -2.24 | 2.08 | 6410.85 | -1.08 | 0.282 | 1.52 | 1.5 | 6490.02 | 1.02 | 0.309 | -6.39 | 2 | 6393.99 | -3.19 | **0.001\*** |
|  | Timebin | 0.34 | 0.68 | 6410.92 | 0.5 | 0.614 | -0.01 | 0.49 | 6490.01 | -0.02 | 0.982 | -6.25 | 0.65 | 6393.58 | -9.57 | **0.000\*** |
|  | Cue × Block | -3.76 | 2.55 | 6410.9 | -1.48 | 0.14 | -3.97 | 1.83 | 6490.02 | -2.16 | 0.03 | 0.09 | 2.45 | 6393.84 | 0.04 | 0.971 |
|  | Cue × Timebin | -4.2 | 0.84 | 6411.23 | -5.02 | **0.000\*** | -2.71 | 0.6 | 6490.01 | -4.52 | **0.000\*** | -0.77 | 0.8 | 6393.82 | -0.96 | 0.337 |
|  | Block × Timebin | 1.22 | 0.96 | 6411.01 | 1.26 | 0.207 | -0.65 | 0.69 | 6490.02 | -0.94 | 0.349 | 2.18 | 0.93 | 6393.95 | 2.35 | **0.019\*** |
|  | Cue × Block × Timebin | 2.34 | 1.18 | 6411.18 | 1.98 | 0.048 | 2.47 | 0.85 | 6490.02 | 2.91 | **0.004\*** | 0.11 | 1.13 | 6393.92 | 0.1 | 0.923 |
| Parietal | Intercept | -6.27 | 2.48 | 71.14 | -2.53 | 0.014 | -8.56 | 1.34 | 89.81 | -6.37 | 0 | 20.78 | 1.82 | 147.48 | 11.45 | 0 |
|  | Cue | 4.38 | 1.94 | 8042.53 | 2.26 | 0.024 | 3.24 | 1.13 | 8364.99 | 2.86 | **0.004\*** | 1.19 | 1.71 | 8136.99 | 0.69 | 0.489 |
|  | Block | -3.13 | 2.23 | 8041.92 | -1.4 | 0.161 | 1.1 | 1.31 | 8364.99 | 0.84 | 0.398 | -5.25 | 1.98 | 8137.42 | -2.66 | **0.008\*** |
|  | Timebin | 2.37 | 0.73 | 8041.9 | 3.23 | **0.001\*** | 2.61 | 0.43 | 8364.99 | 6.09 | **0.000\*** | -7.52 | 0.64 | 8137.03 | -11.69 | **0.000\*** |
|  | Cue × Block | -3.09 | 2.73 | 8042.17 | -1.13 | 0.258 | -3.51 | 1.6 | 8364.99 | -2.19 | 0.029 | -1.18 | 2.42 | 8137.2 | -0.49 | 0.625 |
|  | Cue × Timebin | -3.54 | 0.9 | 8042.61 | -3.94 | **0.000\*** | -2.52 | 0.52 | 8364.99 | -4.82 | **0.000\*** | -0.75 | 0.79 | 8137 | -0.95 | 0.343 |
|  | Block × Timebin | 2.24 | 1.03 | 8041.97 | 2.17 | 0.03 | 0.19 | 0.61 | 8364.99 | 0.31 | 0.754 | 2.9 | 0.91 | 8137.17 | 3.19 | **0.001\*** |
|  | Cue × Block × Timebin | 1.42 | 1.27 | 8042.25 | 1.12 | 0.264 | 1.53 | 0.74 | 8364.99 | 2.07 | 0.039 | 0.06 | 1.11 | 8137.2 | 0.06 | 0.956 |
| **Occipital** | Intercept | -3.24 | 2.99 | 196.13 | -1.09 | 0.279 | -4.23 | 1.69 | 192.31 | -2.5 | 0.013 | 18.91 | 2.5 | 213.08 | 7.56 | 0 |
|  | Cue | 2.77 | 2.98 | 2661.63 | 0.93 | 0.352 | -0.44 | 1.67 | 2745 | -0.27 | 0.791 | 0.06 | 2.52 | 2723.69 | 0.02 | 0.981 |
|  | Block | 0.03 | 3.42 | 2661.2 | 0.01 | 0.992 | -2.26 | 1.93 | 2744.99 | -1.17 | 0.243 | -2.31 | 2.91 | 2724.13 | -0.79 | 0.427 |
|  | Timebin | 1.37 | 1.12 | 2661.08 | 1.22 | 0.221 | 1.62 | 0.63 | 2744.99 | 2.56 | **0.011\*** | -8.01 | 0.95 | 2724.09 | -8.47 | **0.000\*** |
|  | Cue × Block | -5.04 | 4.19 | 2661.26 | -1.2 | 0.23 | 1.02 | 2.37 | 2744.99 | 0.43 | 0.667 | -0.72 | 3.56 | 2723.97 | -0.2 | 0.84 |
|  | Cue × Timebin | -2.2 | 1.38 | 2661.46 | -1.59 | 0.111 | 0.29 | 0.77 | 2745.02 | 0.38 | 0.705 | -0.32 | 1.16 | 2723.68 | -0.28 | 0.781 |
|  | Block × Timebin | 1.53 | 1.59 | 2661.53 | 0.96 | 0.337 | 1.02 | 0.89 | 2744.99 | 1.14 | 0.253 | 2.56 | 1.34 | 2723.97 | 1.91 | 0.056 |
|  | Cue × Block × Timebin | 1.48 | 1.95 | 2661.42 | 0.76 | 0.449 | -0.91 | 1.1 | 2745.01 | -0.83 | 0.408 | -0.58 | 1.64 | 2723.91 | -0.35 | 0.724 |

There was a significant effect of cue on relative alpha band power in central sites (Figure *3*c,e). This effect was due to stronger alpha ERD in central electrodes following pain-related compared to neutral cues (neutral – pain contrast: central, t(6418) = 6.15, *p < .*001). Significant cue × timebin interactions were observed in central and parietal regions (Figure *3*e,g). Breaking the cue effect down by timebin, this effect was driven by cue effects emerging at early and late anticipation intervals. Pain-related cues showed stronger central and parietal ERD compared to neutral cues during early (central, t(6418) = 6.17, *p < .*001; parietal, t(8050) = 5.5, *p < .*001) and late anticipation (central, t(6418) = 7.87, *p < .*001; parietal, t(8050) = 6.92, *p < .*001). Comparatively, during cue presentation, both neutral and pain-related cues were associated with parietal ERD and central ERS in the alpha band (*p >* .05). The effect of block was not significant, nor was there a significant cue × block interaction (all *p >* .05; *Figure 4*a,d).

**Figure 3.**
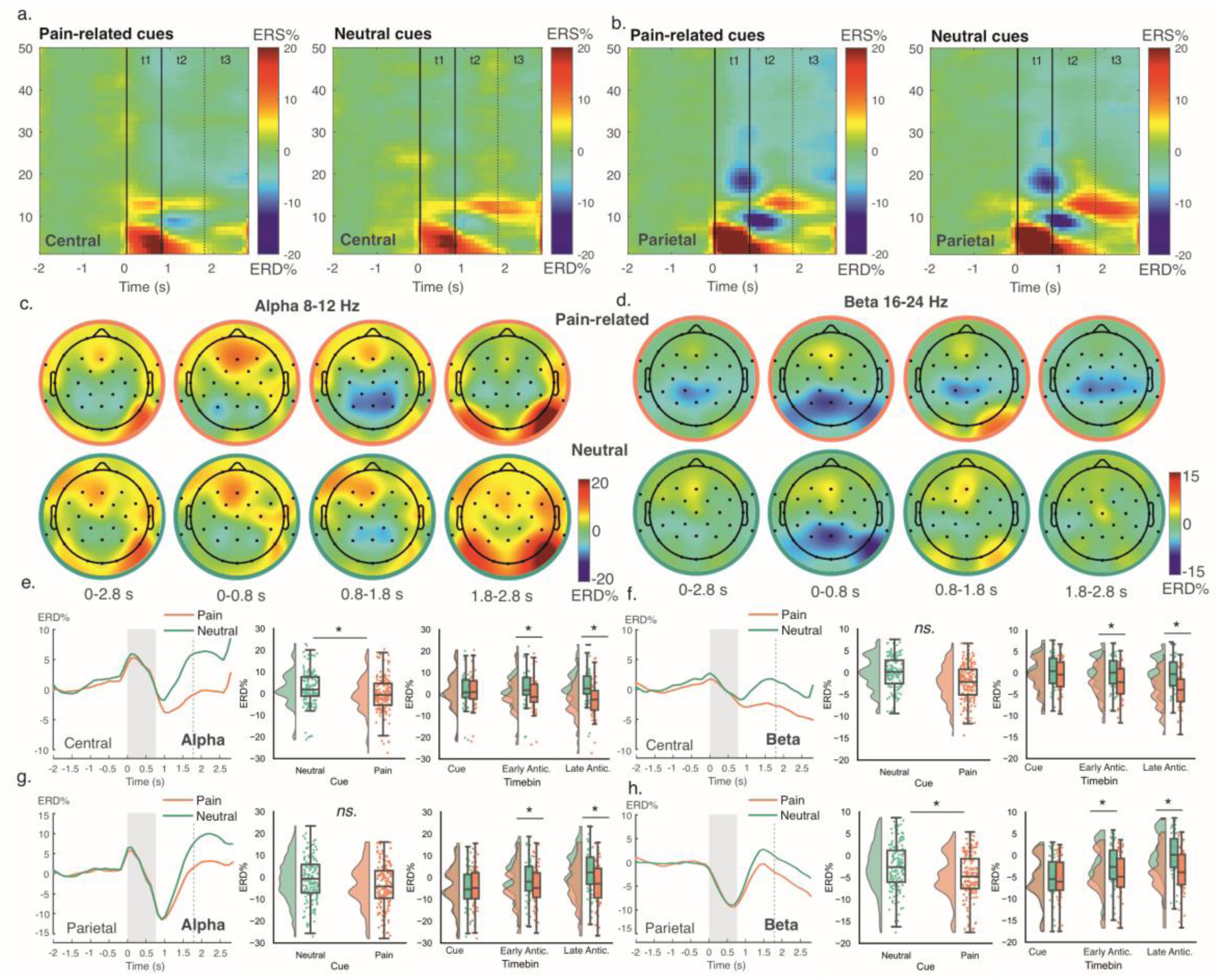
Effect of cue on EEG band power. Time-frequency maps show relative band power changes in central (a) and parietal (b) regions following pain-related versus neutral cues. Topographic maps show ERD/S in alpha (c) and beta (d) frequency bands averaged over the whole trial prior to US onset (0-2.8 s), and during cue presentation (0–0.79 s), early anticipation (0.8–1.79 s) and late anticipation (1.8–2.8 s) intervals. Timecourses show changes during trials in alpha (e, g) and beta (f, h) band power in central and parietal sites for each cue type, averaged over blocks. Shaded regions and dotted lines illustrate CS presentation and separation into early and late anticipation intervals, respectively. Raincloud plots show significant effects in planned comparisons in the corresponding frequency bands, with asterisks indicating significant effects at p < .05. In all plots, negative values indicate event-related desynchronisation (ERD), while positive values indicate event-related synchronisation (ERS).

**Figure 4.**
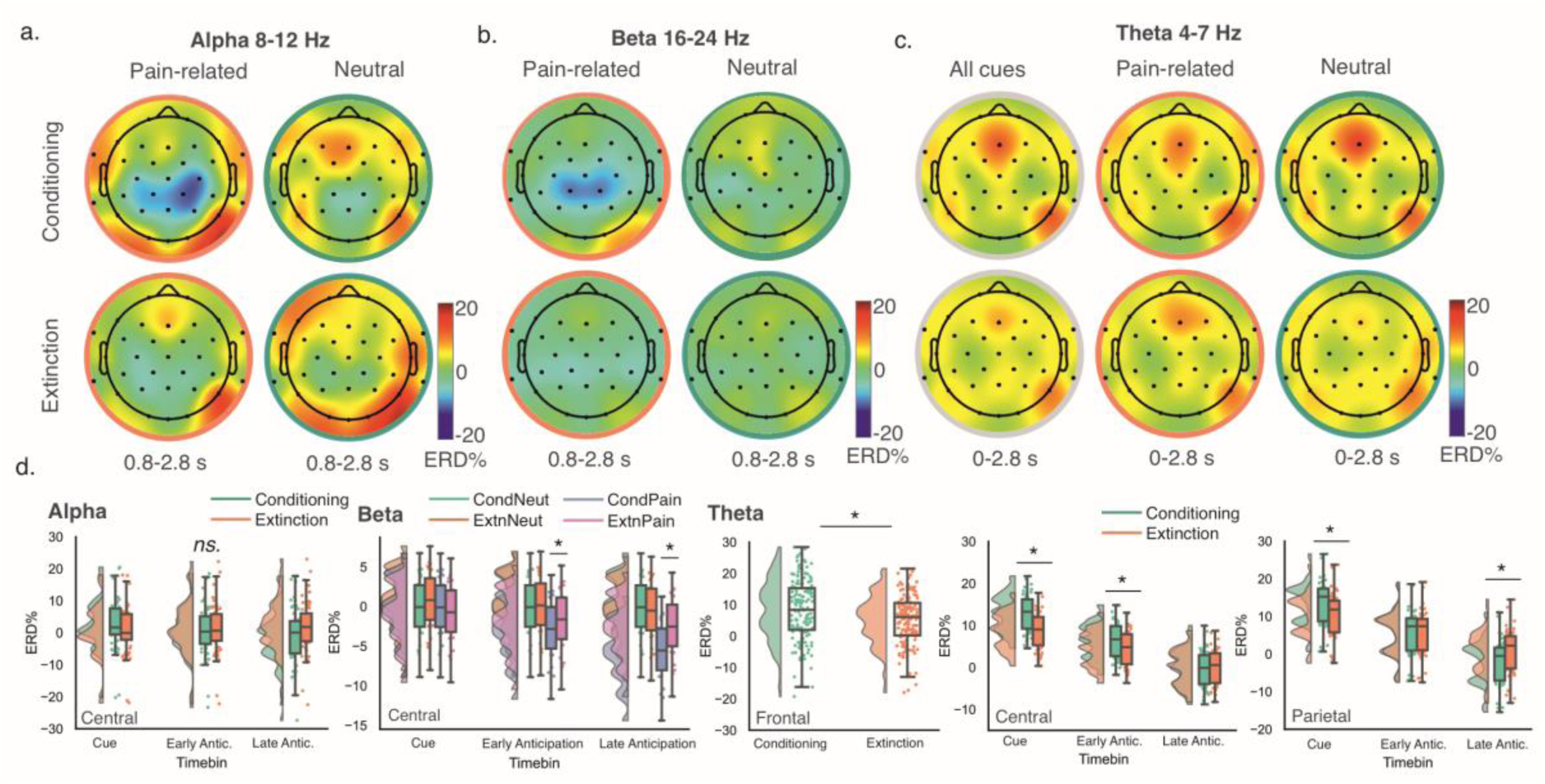
Effect of conditioning block on EEG band power. Topographic maps show ERD/S in alpha (a), beta (b) and theta (c) frequency bands following pain-related versus neutral cues in phase 1 conditioning and phase 2 extinction blocks. Alpha and beta maps are plotted during early and late anticipation intervals (0.8 –2.8 s), while theta maps are plotted throughout the anticipatory period (0–2.8 s), due to significant results in the respective timebins. Raincloud plots (d) show significant effects in planned comparisons in the corresponding bands, with asterisks indicating significant effects at p < .05 after FWE correction. In all plots, negative values indicate event-related desynchronisation (ERD), while positive values indicate event-related synchronisation (ERS).

In the beta band, significant effects of cue on relative band power were found in parietal sites (*Figure 3*d,h) due to stronger ERD for pain-related compared to neutral cues (t(8372) = 6.71, *p < .*001). Significant cue × timebin interactions were observed in central and parietal regions due to divergent effects of cue at early and late anticipation intervals. In parietal sites, both pain-related and neutral cues were associated with beta ERD during cue presentation (*p >* .05), which was sustained for pain cues only (i.e., attenuated for CS-) throughout early and late anticipation intervals (early, t(8372) = 6.71, *p < .*001; late, t(8372) = 7.92, *p < .*001). In central sites (*Figure 3*f), ERD developed throughout the trial for pain cues only, with significant differences based on cue valence appearing during early and late anticipation (early, t(6497) = 6.54, *p < .*001; late, t(6497) = 6.83, *p < .*001).

Cue × timebin × block interactions were found in the beta band over central sites (*Figure 4*). To examine differences in the processing of pain-related (CS+) cues in conditioning compared to extinction blocks, this effect was broken down by cue and timebin. Differences in the processing of pain-related cues between conditioning and extinction blocks were significant during early and late anticipation, with greater ERD for pain-related cues in conditioning blocks (conditioning – extinction pain cues contrast: early, t(6497) = -3.00, *p = .*003; late, t(6497) = -4.78, *p < .*001). No difference between conditioning and extinction blocks was found for neutral cues, nor was any difference found between conditioning and neutral cues during cue presentation (*p >* .05).

The theta band showed a significant effect of block in frontal, central and parietal regions (Figure *4*). In frontal and central regions, this was due to stronger ERS during conditioning versus extinction blocks (frontal, t(2655) = 3.83, *p* < .001; central, t(6401) = 4.06, *p <* .001). The effect of block in parietal regions was not significant in pairwise comparisons after multiple comparison correction (*p >* .05). These effects were qualified by block × timebin interactions in central and parietal regions. In parietal regions, this effect was due to strong ERS during cue presentation which was greater in conditioning versus extinction blocks, and which reduced throughout the trial, manifesting as stronger ERD during late anticipation (cue, t(8145) = 4.03, *p* < .001; late, t(8145) = -4.14, *p <* .001). In central regions, this effect was due to stronger ERS during cue presentation and early anticipation for conditioning versus extinction blocks (cue, t(6401) = 5.61, *p* < .001; early, t(6401) = 4.07, *p <* .001). No block differences were observed during late anticipation (*p >* .05). The effect of cue was not significant, nor was there a significant cue × block interaction (all *p >* .05).

In summary, there was evidence of neural conditioning to the pain cue (CS+) in both alpha and beta bands, and a general effect of block in the theta band.

### 2.3 Computational modelling of trial-by-trial anticipatory learning signals

To investigate neurophysiological learning indices, we examined trial-by-trial ERD/S and pupil diameter following cue onset. We constructed a temporal difference reinforcement learning model fit to trial-by-trial ERD/S and pupil dilation. Linear regression assessed the relationship between the model’s estimated value (*V*) and EEG/pupil measures. Full results are presented in Supplementary Materials.

Across conditioning and extinction blocks, model fit for pupil diameter was statistically significant (F(7,35239) = 155.7, *p* < .001, adjusted R^2^ = .030). Significant interactions between model value and pupil diameter emerged in late anticipation and approach intervals (t = 4.86, *p* < .001, *β* = .06; t = 8.10, *p* < .001, *β* = .10). For EEG metrics, the model fit was also statistically significant (F(35,155972) = 75.33, *p* < .001, adjusted R^2^ = .02). Interactions between learned model value and oscillatory band power were significant in the alpha band over central regions during cue, early, and late intervals (central cue: t = 2.69, *p* = .007, *β* = 3.19; central early: t = -2.46, *p* = .014, *β* = -2.92; central late: t = -5.32, *p* < .001, *β* = -6.33), in parietal alpha band during cue and late intervals (parietal cue: t = 5.05, *p* < .001, *β* = 6.00; parietal late: t = -4.24, *p* < .001, *β* = -5.07); in central beta band during early and late intervals (central early: t = -2.33, *p* =.020, *β* = -2.763; central late: t = -4.13, *p* < .001, *β* = -4.89); and in theta band over frontal regions during cue and early intervals (frontal cue: t = 4.53, *p* < .001, *β* = 5.40; frontal early: t = 5.17, *p* < .001, *β* = 5.40), and over central scalp regions during early intervals (t = 5.17, *p* < .001, *β = 6.14*).

In summary, trial-by-trial analyses echoed results from average condition data showing evidence of neural conditioning, with additional observations during earlier CS-US intervals in alpha and theta bands, which were not observed in the standard LMM analyses.

### 2.4 Effects of Tonic Pain-CS+ congruency

Tonic pain was administered to either laterality in a counterbalanced order during the extinction blocks, allowing us to test our central hypothesis of whether congruency between tonic pain and phasic pain cues modulated behavioural or neural responses. Hence, for these analyses, we looked specifically at the CS+ pain-related cues (i.e., not neutral CS- cues).

Individual LMMs were used to predict changes in pupil diameter and fixations as a function of tonic pain side (congruent versus incongruent with cue laterality) and measurement interval (cue, early anticipation, late anticipation, and rock approach timebins). The main effects of cue congruency and timebin were not significant for pupil diameter, nor were there any significant cue × timebin interactions (all *p* > .05; **Table 1**).

For neural data, grand average time-frequency and topographic plots for pain-related trials during the extinction block are shown in Figure *5*. Individual LMMs were fit for each frequency band and electrode cluster, with fixed effects of tonic pain side and measurement interval (cue, early, and late anticipation) (**Table 3**). A significant effect of tonic pain congruency was found in frontal electrodes in the theta band. This was due to a reduced focus of ERS across frontocentral sites for congruent vs. incongruent cues, which reached significance following FWE-correction in the frontal regions (t(2134) = 2.62, *p =* .008; Figure ***5***a,e). No statistically significant tonic pain congruency effects were observed in alpha or beta bands (*p* > .05).

**Figure 5.**
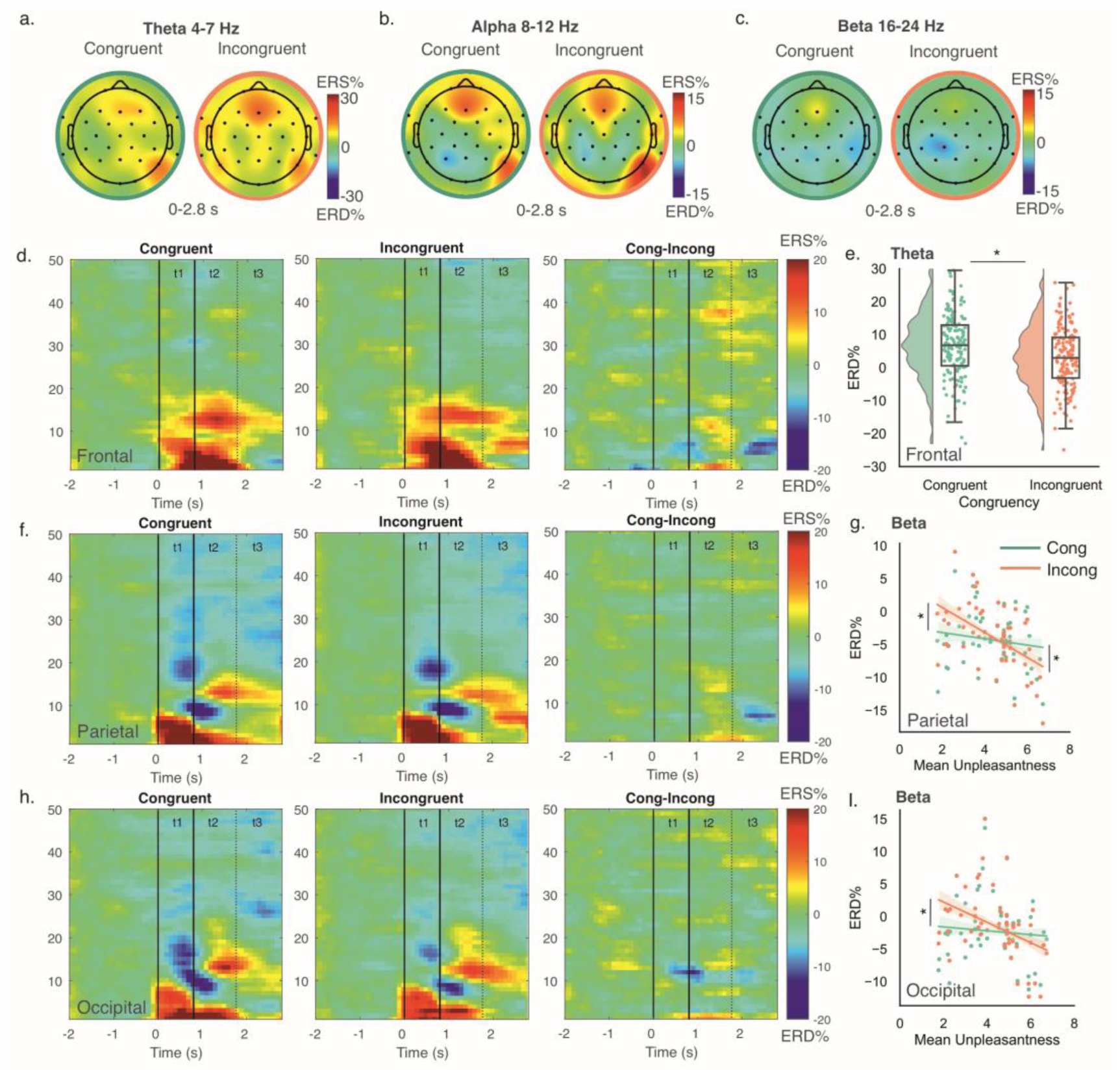
Effect of tonic pain on EEG band power during extinction. Topographic maps show ERD/S during tonic pain on sites congruent or incongruent to the cued location in theta (a), alpha (b) and beta (c) frequency bands throughout the anticipatory period (0–2.8 s). Time-frequency plots show relative band power changes for congruent, incongruent, and congruent minus incongruent trials in frontal (d), parietal (f) and occipital (h) electrode sites. Plots (e, g, i) show significant effects of tonic-phasic pain congruency in planned comparisons in the theta band over frontal sites, and in the beta band over parietal and occipital sites when adjusting for mean unpleasantness ratings of tonic pain. In all plots, negative values indicate event-related desynchronisation (ERD), while positive values indicate event-related synchronisation (ERS). Asterisks indicate significant effects at p < .05 after FWE correction.

**Table 3.** EEG fixed effects for tonic pain – cue congruency and timebin. Asterisks denote effects that survived FWE correction (*p* < .05). Default (treatment) contrast coding was used for measurement timebin (reference: cue) and tonic pain side (reference: incongruent).

| Cluster | Predictor | Band |  |  |  |  |  |  |  |  |  |  |  |  |  |  |
| --- | --- | --- | --- | --- | --- | --- | --- | --- | --- | --- | --- | --- | --- | --- | --- | --- |
|  |  | Alpha |  |  |  |  | Beta |  |  |  |  | Theta |  |  |  |  |
|  |  | *b* | *SE* | *df* | *t value* | *p* | *b* | *SE* | *df* | *t value* | *p* | *b* | *SE* | *df* | *t value* | *p* |
| Frontal | Intercept | 7.47 | 2.06 | 80.21 | 3.63 | 0.000 | 2.99 | 1.25 | 98.05 | 2.39 | 0.019 | 19.53 | 2.15 | 99.45 | 9.06 | 0.000 |
|  | Congruency | 1.75 | 1.04 | 861.63 | 1.68 | 0.094 | 0.54 | 0.66 | 884.00 | 0.82 | 0.413 | -3.73 | 1.16 | 842.80 | -3.23 | 0.001* |
|  | Timebin | -2.02 | 0.63 | 860.91 | -3.18 | 0.002 | -1.29 | 0.41 | 884.00 | -3.17 | 0.002 | -6.45 | 0.70 | 837.02 | -9.19 | 0.000 |
| Central | Intercept | 0.12 | 1.83 | 45.50 | 0.06 | 0.949 | -0.24 | 1.00 | 73.42 | -0.24 | 0.815 | 14.12 | 1.35 | 83.50 | 10.45 | 0.000 |
|  | Congruency | 0.96 | 0.68 | 2118.23 | 1.41 | 0.158 | 0.56 | 0.48 | 2131.01 | 1.16 | 0.245 | -0.50 | 0.68 | 2088.13 | -0.74 | 0.459 |
|  | Timebin | -0.29 | 0.42 | 2117.56 | -0.68 | 0.494 | -0.90 | 0.30 | 2131.01 | -3.03 | 0.002 | -4.82 | 0.42 | 2086.38 | -11.57 | 0.000 |
| Parietal | Intercept | -8.33 | 2.14 | 41.81 | -3.90 | 0.000 | -7.55 | 1.04 | 53.84 | -7.25 | 0.000 | 15.13 | 1.44 | 72.91 | 10.54 | 0.000 |
|  | Congruency | 0.51 | 0.75 | 2658.56 | 0.68 | 0.497 | -0.34 | 0.44 | 2754.01 | -0.78 | 0.438 | -0.49 | 0.68 | 2685.63 | -0.71 | 0.475 |
|  | Timebin | 2.34 | 0.46 | 2658.48 | 5.09 | 0.000 | 1.80 | 0.27 | 2754.00 | 6.76 | 0.000 | -5.06 | 0.42 | 2684.55 | -12.05 | 0.000 |
| Occipital | Intercept | -5.16 | 2.11 | 101.85 | -2.45 | 0.016 | -5.49 | 1.30 | 71.25 | -4.24 | 0.000 | 16.20 | 1.97 | 71.11 | 8.22 | 0.000 |
|  | Congruency | 0.14 | 1.13 | 863.79 | 0.12 | 0.902 | -0.85 | 0.62 | 884.00 | -1.38 | 0.169 | -0.66 | 0.94 | 872.86 | -0.71 | 0.478 |
|  | Timebin | 1.80 | 0.69 | 862.91 | 2.59 | 0.010 | 2.03 | 0.38 | 884.00 | 5.34 | 0.000 | -6.42 | 0.57 | 873.41 | -11.17 | 0.000 |

Next, we performed exploratory covariate analyses to identify any additional modulation of congruency effects. Age, sex, STAI trait scores and mean self-reported tonic pain ratings (intensity and unpleasantness) were added to individual LMMs to examine the effect of covariates on changes in pupil diameter and EEG power spectra. No covariate significantly predicted the change in pupil diameter during anticipation, nor did the addition of a covariate in the model result in a significant congruency effect (*p* > .05).

For EEG data, including covariates in LMMs resulted in significant effects not evident with the condition structure alone (Supplementary Materials; Figure *5*). Following FWE corrections, these results were restricted to the beta band. Accounting for tonic pain ratings resulted in significant main effects of congruency in parietal and occipital sites (Figure *5*g,i), with stronger beta-band ERD for congruent vs. incongruent cues. This effect was not significant in pairwise comparisons (*p* > .05) but was qualified by significant cue × rating interactions. Examination of the interaction showed an inverse relationship between ERD/S and mean unpleasantness in parietal and occipital beta band power, specifically for incongruent cues. Specifically, participants with the lowest unpleasantness ratings showed attenuated parietal and occipital ERD (/greater ERS) for incongruent vs. congruent cues (parietal, t(2756) = 3.89, *p <* .001; occipital, t(886) = 3.01, *p =* .003). Conversely, participants reporting the highest tonic pain unpleasantness showed stronger parietal ERD for incongruent cues (t(2756) = -3.09, *p =* .002; occipital, *p* > .05). In other words, higher tonic pain ratings were associated with weaker parietal and occipital beta ERD for congruent compared to incongruent cues. No significant differences were found at mean unpleasantness ratings (*p* > .05).

### 2.5 Computational modelling of trial-by-trial Tonic Pain-CS+ congruency

Individual linear regression analyses assessed the relationship between EEG/pupil measures and learned model values during extinction blocks with tonic pain. For pupil diameter, the overall fit of the model was statistically significant (F(3,13917) = 9.84, *p* < .001, adjusted R^2^ = .002), but no interactions between learned values and congruency were observed. For EEG data, the overall fit of the model was statistically significant (F(3,61803) = 3.22, *p* = .022, adjusted R^2^ = .00), showing a steeper positive relationship between *V* and single-trial ERD/S for congruent versus incongruent cues (t = 3.07, *p* = .002, *β* = -5.55). When examining this change within frequency bands, there was no significant interaction between learned values, congruency and band power after FDR correction, although a trend towards a significant effect in the theta band was noted (see Supplementary Materials).

In summary, under tonic pain, EEG oscillations at the average condition level showed reduced midfrontal theta ERS for congruent vs. incongruent cues; with this signal partially captured by trial-by-trial changes in cue valuation.

## 3. Discussion

We investigated how tonic pain influences neural representations of phasic pain predictions using a two-phase integrated EEG-VR revaluation task. Participants were first conditioned to predict phasic pain to the left or right arm. EEG data characterised the neural conditioned response, with greater central and parietal alpha and beta ERD associated with pain predictions. In the subsequent extinction phase when tonic pain was applied, there was reduced midfrontal theta ERS when the laterality of tonic pain matched (was congruent with) the laterality prediction of phasic pain. When accounting for the perceived magnitude of tonic pain ratings, tonic-phasic congruency also identified greater parietal and occipital beta ERD, in that congruence boosted pain predictions manifesting in greater posterior beta ERD correlating with the unpleasantness of tonic pain. Taken together, the data provide evidence of the presence of an internal representation of the specific cue-pain memory that is modulated by tonic pain.

This study was motivated by the concept that tonic and phasic pain serve different functions: whilst phasic pain acts as a rapid alarm and teaching signal for potential harm, tonic pain provides an ongoing homeostatic signal of presumptive actual harm ^12^ ^2,3,13^. This theory derives from the assertion that tissue damage will result in physiological vulnerability of that area of the body, and a function of tonic pain is to drive additional protective behaviours to help reduce further harm to the area. Well-known components of this are peripheral and central pain sensitization, whereby hyperalgesia and allodynia amplify and transform the painfulness of new phasic stimuli in the area of damage ^14^. However, the data here support the idea that the brain goes further, by modifying the *prediction* of stimuli to that region. This provides extra protective mechanisms without the need to directly experience new stimuli to the vulnerable area, applicable in any setting where pain can be predicted.

Thus, defensive actions can in principle be taken *before* a new potentially harmful stimulus occurs, illustrating the critical time advantage of using learned predictions to optimise behaviour. This capacity to revalue predictions (i.e., memories) according to global physiological priorities is a hallmark of homeostatic states that is well recognised for reward: e.g., where selective satiation modifies food predictions for specific food categories ^15,16^. However, whilst homeostatic theories of pain are widely proposed ^2,12,17^, this fundamental aspect of pain processing had not previously been directly tested.

Modification of phasic pain predictions by tonic pain inherently implies that the brain forms an internal representation of the pain: a memory of the cue-pain association which can be subsequently modified. Importantly, this representation must include a topographic representation of the pain predicted (with evidence of anticipatory topographically organised pain predictions manifesting in lateralised motor adaptations at the threat of pain ^18^), so that the prediction of the phasic pain site can be internally compared against the site of sustained pain. In classical learning theory, the modulation of predictions by homeostatic states is taken a cardinal feature of internal representations of ‘models’, as it shows that predictions are not merely cue-driven processes but hold information that specifically relate to the outcome.

Thus, revaluation/devaluation paradigms have played an important role in animal learning studies, at the heart of the distinction between ‘model-based’ and ‘model-free’ cognition and decision making ^19^. The data here suggest that pain learning engenders model-based processes, in which pain-cue associations are explicitly stored in the brain, over and above simple elicitation of cue-driven or reflexive responses. This adds to other evidence for model-based representations for phasic pain, including in the distinction between trace and delay conditioning ^20^, and in multi-step decision-making ^21^.

Neural responses associated with pain predictions can be related to previous studies. During conditioning, pain-predictive cues elicited robust central-parietal alpha and beta suppression following cue offset. Diminished alpha band power has been linked to the salience or valence of conditioned stimuli ^9,22^ and sustained attention ^23,24^, while beta oscillations have been linked to motor preparation, sensory perception, and selective attention ^25–29^. Consistent with previous aversive conditioning literature ^9^, conditioned responses (CS+ compared to CS-cues) were observed and sustained throughout early and late CS-US intervals.

The key analysis was in neural activity linked to congruence, as this reflects the computation of enhanced threat/danger that occurs when a phasic insult is predicted to an area of presumed existing damage. We identified diminished frontal theta synchronisation during congruent versus incongruent cues. Theta synchronisation was not identified in simple conditioning contrasts of cue valence (CS+ compared to CS-), with generally reduced ERS during extinction versus conditioning blocks. This counters the notion of a simple value response, but supports the idea that the brain performs the underlying computations to recognise the significance of congruence. A potentially useful association here is with theories of control contexts, in which diminished theta is associated with greater utilisation of Pavlovian control mechanisms, particularly when such responses are adaptive ^30^. A shift towards Pavlovian responses is behaviourally appropriate in this context, where pain prediction could reflect an enhanced protective response.

Incorporating experimental subject variability in the perceived experience of tonic pain allowed us to further probe EEG correlates of congruency. Higher tonic pain ratings were associated with weaker parietal and occipital beta ERD for congruent cues compared to incongruent cues. Enhanced beta ERD was also identified as a basic conditioned response to CS+ cues in conditioning phases. Therefore, this does not seem to represent a simple value response, which would be enhanced with congruent cues. Instead, it may represent a reduced inhibitory response such as endogenous pain modulation, in that a weaker ability to exert pain inhibition during congruence may lead to enhanced pain experience. Alternatively, it could reflect additional active processes related to attending to multiple locations: an existing area of presumed damage (tonic pain) at one site alongside threat of damage (phasic pain prediction) at another site. Aligning this with the broader EEG literature, increased beta ERD could relate to differences in motor readiness ^31,32^ and sensorimotor preparation ^28,29,33^ to process topographically conflicting cues under heightened aversiveness. This is in accordance with evidence that pain disrupts attentional processes, particularly relating to switching and orienting attention ^34^, as well as deficits in attention in chronic pain ^35^. It also illustrates a key role of individual experiences in shaping the neural dynamics of pain valuation. More broadly, however, this result provides further evidence that the brain detects and recognises the key distinction between congruent and incongruent pain predictions.

Trial-by-trial modelling illustrated that changes in pupil diameter and EEG power were predicted by learned cue values, reflecting dynamic updates in pain anticipation. Notably, significant correlations emerged between learned values and pupil dilation during late anticipation and approach trial intervals. Echoing findings from averaged EEG, correlations between learned cue values and single-trial EEG were observed in central alpha and beta (early/late intervals) and parietal alpha (late interval). Unique to modelling analyses, central and parietal alpha power also correlated with learned values during cue presentation, and frontal and central theta correlated positively with learned values during cue and early anticipation intervals. Furthermore, under tonic pain, EEG oscillations showed a generally stronger negative correlation with learned values for congruent versus incongruent cues. Notably, we observed a trend towards this effect specifically in midfrontal theta band power, similar to previous literature on Pavlovian influences on action in response inhibition tasks ^30^ ; although this effect was not significant after multiple comparison correction. Further work should further investigate the potential role of theta oscillations in mediating cue valuation processes.

Several caveats should be noted. First, this study did not include a clear behavioural demonstration that neural evidence of revaluation manifests in enhanced defensive actions. Physiological measures reflected heightened arousal for pain-predictive cues but were not modulated by tonic pain laterality. Pupil dilation increased during later anticipation intervals, and fixations were more frequent during pain-cue approach interval, reflecting heightened arousal for pain-predictive cues, consistent with previous work ^40–42^. Ideally, future experiments to test behavioural effects of congruence should extend to an instrumental context, allowing subjects to make choices to avoid Pavlovian cues that predicted phasic pain to the tonic pain area (such as conditioned suppression or conditioned negative reinforcement).

A second caveat is that the neural data does not distinguish activity related to the actual encoding of the cue-pain memory from activity related to behavioural processes that arise from it; therefore, we cannot draw firm conclusions about the nature of representations of the encoded pain memory itself. Despite this, we can use the neural data to infer that some neural encoding of pain memories must be represented in some specific manner in the brain.

Third, devaluation and revaluation paradigms necessarily test behaviour during extinction, to avoid contaminating the revalued conditioned response with new learning of directly experienced outcomes (i.e. the phasic pain). Hence the Pavlovian associative memory is expected to decline during the critical test block. This motivates the computational analysis, which captures the trial-by-trial dynamics of how values are expected to diminish throughout the extinction process, but is dependent on how well the model captures the learning process.

In summary, the results define a novel process by which ongoing tonic pain reshapes neural activity associated with predictions of phasic pain in a topographically specific manner. This provides evidence for a specific but previously untested prediction of the broader homeostatic pain hypothesis, illustrating the multifaceted nature of pain acting at different temporal profiles, and the complexity of adaptive physiological pain behaviour.

## 4. Methods

### 4.1 Participants

Thirty-one pain-free participants (14 women) with no neurological conditions were recruited from the community and a pool of university staff and students. Five subjects were excluded during data collection: three due to technical issues during the recordings, and two due to not adhering to the study protocol. The final sample included 26 participants (13 women; 24 right-handed, 1 ambidextrous) with a mean age of 26 ± 4.9 years (mean ± SD). The procedure was approved by the Research Ethics Committee of the University of Oxford, and all participants gave written informed consent at the start of the experiment following the Declaration of Helsinki. Participants were reimbursed with £60 on completion of the study.

### 4.2 Procedure

Experimental procedures were carried out in a single 3-hour session at the Oxford Institute of Biomedical Engineering. Participants were seated in a comfortable chair while EEG electrodes and the VR headset were applied. Participants were asked to hold the VR handsets with arms shoulder width apart, resting on an armrest raised at a 45-degree angle from the desk.

Participants viewed a realistic rendering of a simulated forest environment (see Figure 1 for full details). The experiment consisted of 4 blocks, divided into 2 interspersed conditioning and extinction phases. During conditioning blocks, left and right looming objects were followed by electrical pain on the corresponding forearm with a 75% shock contingency (pain-related cues; determined automatically at the start of the experiment). No pain was delivered following central looming objects (neutral cues). A self-timed break followed every 24 trials, with 24 trials per cue (72 trials for each block; 144 trials overall).

During extinction blocks, tonic pain was applied using a pressure cuff. During these trials, rocks travelled in the cued direction but disappeared 0.1 s early. No shocks were delivered. There were 108 trials for each block (216 trials overall): 36 for each cue type, with a self-timed break of at least 1 minute for every 12 trials where the pressure cuff was deflated.

Throughout the task, trials were ordered pseudorandomly by shuffling the order of 6 trials (2 repetitions of each cue type) and repeating for the total trial number. During each break, participants were asked to rate the intensity and unpleasantness of electrical (conditioning blocks) or pressure (extinction blocks) pain stimuli on a scale of 0 (not painful/unpleasant), 3 (just painful/unpleasant) to 10 (maximum pain/unpleasantness) for each site. At the end of the experiment, participants were asked to complete a brief questionnaire regarding demographic characteristics and the State-Trait Anxiety Inventory.

### 4.3 Pain Stimuli

#### 4.3.1 Phasic Electrical Pain Stimuli

Electrical pain stimulation was applied to the skin of the forearm using EPS-P10 electrodes (EPS-P10, MRC Systems GmbH, Germany) connected to a constant current stimulator (Digitimer DS7a, Digitimer Ltd, UK). Electrical stimulation intensity was set for each participant as a multiple of electrical detection threshold (EDT). EDT was determined for each forearm at the start of the experiment using the method of limits, where single electrical stimuli were delivered in descending and ascending steps of 0.002 mA to establish the lowest perceptible threshold. Participants rated single electrical stimuli at a multiple of EDT (10–30×) until an intensity of 5/10 on a scale of not painful (0), just painful (3), to most pain imaginable (10) was reached, following previous studies ^36–38^. During the main experiment, stimuli consisted of 2 rapidly succeeding pulses (frequency 1 Hz, pulse width 1 ms, duration 1 ms) at the test stimulation intensity. If participants reported that the stimuli were no longer painful during breaks, intensity was increased; likewise, if the stimuli became intolerable, intensity was decreased.

#### 4.3.2 Tonic Pressure Pain Stimuli

Tonic pain was delivered to the left and right upper arms using a manual pneumatic blood pressure cuff. Nociceptive responses elicited by pressure stimuli have been demonstrated to originate from deep tissue muscle strain rather than superficial cutaneous or ischemic sensations ^39,40^. Stimulation intensity was determined at the start of the experiment using a staircase procedure, where the pressure was slowly increased until participants reported a ‘just painful’ sensation (3/10).

### 4.4 Virtual-Reality and EEG Equipment

The experiment was designed in Unity and viewed with a HTV Vive Pro Eye headset, connected to SteamVR using an Alienware PC. Eye-tracking data was collected using the headset following calibration at the start of the experiment.

#### 4.4.1 Physiological data analysis

Physiological VR data were processed in Jupyter using Python 3. Trial data were extracted from continuous files for each participant for the whole trial period, starting -1 s prior to cue presentation to 3.3 s after cue onset. Data were interpolated to account for slight variations in sampling rate throughout the trials, to a total of 3300 datapoints per trial.

Pupil diameter for left and right pupils was extracted from trial data. To account for ocular artefacts such as blinks, data points exceeding 2 SDs from the trial mean were identified. Where 150 consecutive extreme data points were found (0.15 s), datapoints were removed and linearly interpolated following previous studies ^41–43^ and according to minimum blink duration ^44^. Trials containing >= 25% interpolated data were removed from further analysis (0 trials). Cleaned data from left and right eyes were individually baseline corrected (-250 to -750 ms) by subtracting the average of the baseline period from the trial period ^41,43^, and subsequently averaged.

The number of fixations on the looming rock were extracted from the last 500 ms of the trial and averaged for each condition. Gaze direction and hand position were extracted from trial data. No significant changes in gaze direction or mean displacement of hand position were observed throughout (full methods and results in Supplementary Materials) (*p* > .05).

#### 4.4.2 EEG acquisition

Whole-scalp EEG was continuously recorded using a 32-channel system (BrainProducts GmbH, Munich, Germany). Actively-shielding Ag-AgCl electrodes were mounted on an electrode cap (actiCap snap, BrainProducts) according to the International 10– 20 system ^45^. The cap was aligned with three anatomical landmarks of two preauricular points and the nasion. Electrolyte gel was applied to achieve electrode-to-skin impedances below 25 kΩ throughout the experiment. A recording band-pass filter was set at 0.001–131 Hz. The sampling rate was 500 Hz, corresponding to a sampling interval of 2000 µs. Electrode FCz was used as a reference electrode, and electrode FPz was used as the ground electrode. EEG average reference was applied, and signals were digitised with a LiveAmp amplifier, connected to BrainVision Recorder 1.25 running on an Alienware PC. Due to a technical issue with the Bluetooth connection between the LiveAmp and the PC, 0.64 ± 1.86 trials were lost for each participant and block.

#### 4.4.3 Spectral analysis of EEG signals

EEG data were processed using EEGLab ^46^. The full processing pipeline is presented in Supplementary Materials. Briefly, continuous EEG data were split into 8 s epochs (-2.5– 5.5 s around cue onset). Data were re-referenced to the common average ^47^ and filtered from 1–70 Hz, with a notch filter from 48–52 Hz. Data for all 4 conditions were subsequently merged, resulting in one datafile per participant. Oculomotor and muscular artefacts were removed from the data using independent component analysis ^48^. Data were epoched from - 2.5–3.5 s around cue onset. Electrode channels with large artefacts were interpolated to a maximum of 10% of all electrodes. Epochs containing extreme data were excluded using a semi-automated method based on epoch amplitude and SD. Data were transformed into current source density using a Laplacian Spherical Spline interpolation method ^48,49^.^51^ Power spectra were computed using a discrete Fourier time-frequency transformation (1–70 Hz, frequency resolution 1 Hz). Relative power was evaluated using the classical ERD transformation,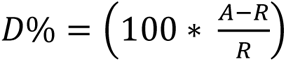, where *D* represents the percentage power change during epochs following cue onset (*A)* relative to the reference period (*R,* -2 to -1 s) ^50^. Positive and negative *D* values correspond to power increases (ERS) ^51^ and decreases (ERD) ^52,53^, respectively.

Relative band power in alpha (8–12 Hz), beta (16–24 Hz) and theta (4–7 Hz) frequency bands was extracted from all 32 active scalp electrodes in 3 timebins: 0–0.79 s, 0.8–1.79 s and 1.8–2.8 s (cue, early, and late anticipation, respectively). Electrodes were clustered into 4 spatially adjacent regions: frontal (Fp1, Fp2, Fz), central (FC5, FC1, FC2, FC6, C3, Cz, C4), parietal (CP5, CP1, CP2, CP6, P7, P3, Pz, P4, P8) and occipital (O1, Oz, O2), based on previous literature showing prominent band power changes during spatial attention, aversive learning and pain anticipation ^10,29,54,55^. No hemisphere lateralisation effects were observed (see Supplementary Materials).

### 4.5 Statistical Analysis

#### 4.5.1 Self-reported measures

For each tonic and phasic pain, two 2×2×3 ANOVAs (left vs. right side; within-block rating time; block repetition (conditioning/extinction block 1 or 2) assessed changes in pain intensity and pain unpleasantness using SPSS v. 29 (IBM Inc., USA). For significant main effects, Bonferroni-corrected pairwise comparisons are reported. Repeated-measures ANOVAs and corresponding pairwise comparisons are reported for interactions.

#### 4.5.2 EEG and Physiological Data

ERD/S and physiological data were entered into individual LMMs in RStudio (Version 2023.12.1.402; ^56^). LMMs were computed with lme4 ^57^ and lmerTest packages ^58^ using Satterthwaite approximation for degrees of freedom. Prior to LMMs, data were checked for extreme values to minimise the influence of outliers on model estimates. Extreme values were identified using z-scores, with values beyond ±3 standard deviations from the mean removed.

Separate LMMs assessed: i) changes as a function of block (conditioning and extinction) and cue (neutral, pain-related), and ii) changes as a function of tonic pain side. Both LMMs additionally included measurement interval (cue, early, late, and approach timebins). The approach timebin was not included in EEG analyses due to the presence of moving stimuli. For EEG data, LMMs were constructed individually for alpha (8–12 Hz), beta (16–24 Hz) and theta (4–7 Hz) frequency bands and electrode clusters, as it was expected that relative band power would differ between frequency bands and topographically over the scalp.

For all LMMs, participants and block repetition were modelled as hierarchical random effects. Models were estimated using ML and nloptwrap optimizer. Fixed effects structures were determined stepwise by including interaction terms or predictors in the model. Predictors and interaction terms which did not significantly change the variance explained were removed to avoid overparameterisation. Bayesian Information Criterion and Akaike Information Criterion were compared for each model to find the simplest model that explained the data.

Exploratory analyses investigated the influence of covariates on the effect of tonic pain-cue congruency. Covariates of interest were age, sex, STAI trait score, mean intensity and mean unpleasantness of tonic pain. Each predictor was added to a covariate-only model. Covariates that significantly predicted change in the measured variable were included in LMMs with the factor of tonic pain side.

*p*-values for all main effects of interest and interactions were adjusted using family-wise error correction (FWE). Significant main effects which surpassed FWE-correction were followed up with pairwise comparisons using the emmeans package. Significant interactions were followed up interaction contrasts for estimated means. All contrasts were adjusted for multiple comparisons using the Šidák correction.

#### 4.5.3 Computational model analysis of trial-by-trial anticipatory learning

Computational learning models were used to capture pain learning in ERD/S and pupil dilation at the trial level. We constructed a standard temporal difference reinforcement learning model, as previously implemented for human pain conditioning ^59,60^. The model is a simple, “real-time” instantiation of the Rescorla-Wagner model ^61^, where the value *V* of trial *n* + 1 for a cue *j* is updated based on the value of current trial *n* and the prediction error, the difference between current value *Vj*, and outcome stimulus value *R* at trial *n*, weighted by a constant learning rate α:

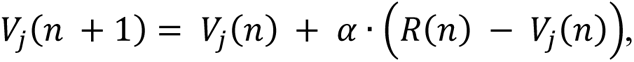

where the learning rate α (0 ≤ α ≤ 1) is a free parameter. Regression analyses (ordinary least-squares) were implemented separately for EEG (on a compound variable of frequency, time and cluster) and physiological data to assess the relationship between learned model values and measured responses during conditioning and extinction blocks, and during extinction blocks based on tonic pain laterality. *p*-values for all effects were adjusted using FWE-correction.

## Data availability

Source data is available at https://osf.io/w25k4/. Analysis code is available at https://github.com/dhewitt16/PainAnticipationVRStudy.

## Acknowledgements

This study was funded by the Wellcome Trust (214251/Z/18/Z) and supported by the NIHR Oxford Health Biomedical Research Centre (NIHR203316). The views expressed are those of the author(s) and not necessarily those of the NIHR or the Department of Health and Social Care. The Wellcome Centre for Integrative Neuroimaging is supported by core funding from the Wellcome Trust (203139/Z/16/Z and 203139/A/16/Z). For the purpose of open access, the author has applied a CC BY public copyright licence to any Author Accepted Manuscript version arising from this submission. The authors would like to thank Dr Charlotte Krahé for sharing their expertise in linear mixed-effects models, and Pranav Mahajan for reviewing the manuscript and advising on reinforcement learning algorithms.

## Author information

**Declaration of Interest:** None of the authors have potential conflicts of interest to disclose. Author roles: DH contributed to conceptualisation, methodology, software, investigation, formal analysis, writing - original draft, writing - review & editing; ST contributed to software, investigation, writing – review & editing; SS contributed to investigation, formal analysis, writing – review & editing; BS contributed to conceptualisation, funding acquisition, supervision, writing - original draft, writing - review & editing.

