## Supplementary Materials for "Tonic pain revalues associative memories of phasic pain"

#### A – Pain ratings

**Table 1. Self-reported ratings and stimulus thresholds prior to the experiment.**

| | | Detection Intensity | Test Intensity | Rating Mean (0-10) |
|---|---|---|---|---|
| Phasic pain | Left arm | *.08 ± .05 mA* | *1.47 ± 1.2 mA* | ***4.85** ± 1.05* |
| | Right arm | *.15 ± .13 mA* | *2.12 ± 1.91 mA* | ***4.42** ± 1.27* |
| Tonic pain | Left arm | *105.00 ± 58.4 bar* | *178.85 ± 76.8 mA* | ***3.38** ± .98* |
| | Right arm | *93.08 ± 52.82 bar* | *169.62 ± 65.51 mA* | ***3.27** ± .96* |

**Phasic (electrical) pain:** Pain intensity ratings showed a significant main effect of block (F(1,25) =4.31, $p$ = .048) due to lower pain intensity ratings in the second versus the first conditioning block ($p$ = .048). The time-by-block interaction was significant (F(2,50) =6.61, $p$ = .003) due to decreased pain intensity ratings within the second conditioning block (F(2,50) =5.9, $p$ = .005), but not the first (F(2,50) =1.33, $p$ = .273). This effect disappeared after controlling for electrical pain intensity on left and right arms (F=2, 46) = 2.31, $p$ = .111). No significant effects of side, time within block, or other interactions were observed.

Pain unpleasantness ratings significantly decreased within- (F(2,50) =4.27, $p$ = .019) and between-blocks (F(2,25) =19.08, $p$ < .001). This was due to a reduction in unpleasantness ratings as the experiment progressed within blocks (first-last block: $p$ = .036). Reductions between blocks remained significant after accounting for stimulation intensity (F(2,23) = .10.98, $p$ = .003), while within-block changes were no longer significant (F(2,46) = .32, $p$ = .720). No other significant effects were found.

**Tonic (pressure) pain:** Pain intensity and unpleasantness ratings decreased significantly within-blocks (intensity, F(9,189) = 12.74, $p$ < .001; unpleasantness, F(9,189) = 12.3, $p$ < .001), following a linear trend (intensity, F(1,21) =18.57, $p$ < .001; unpleasantness, F(1,21) =13.07, $p$ = .002). The effect was significant only after the first 2 minutes (time 0 – time 1: intensity, $p$ = .007; unpleasantness, $p$ = .001). Time-by-cuff site effects were significant (intensity, F(9,189) =2.07, $p$ = .034; unpleasantness, F(9,189) =2.16, $p$ = .027), with greater reductions in ratings over time when the cuff was on the right arm (all $p$ < .05) vs. the left arm (all $p$ >.05, except for time 1 vs. 4 for pain unpleasantness). Block-by-time effects showed stronger reductions in intensity for the first versus the second extinction block (intensity, F(9,189) =2.67, $p$ = .006; unpleasantness, F(9,189) =2.53, $p$ = .009). For unpleasantness, the decrease in pain ratings between time 1 and all subsequent timepoints was significant for both blocks (all $p$ < .005). The effect of cuff site was not significant.

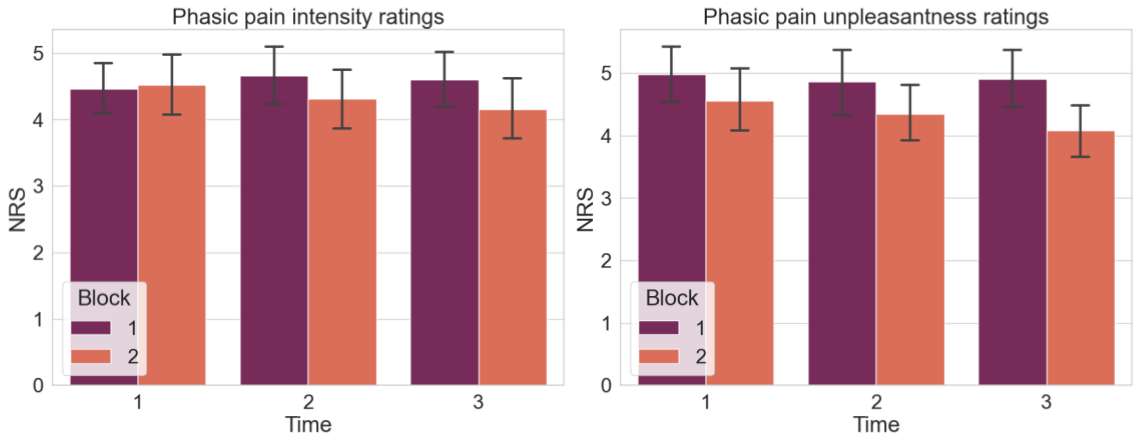


Figure 1. Plot of electrical pain ratings (averaged over left and right sides) during conditioning blocks 1 and 2. Ratings were taken after 24 (Time 1), 48 (2) and 72 trials (3).

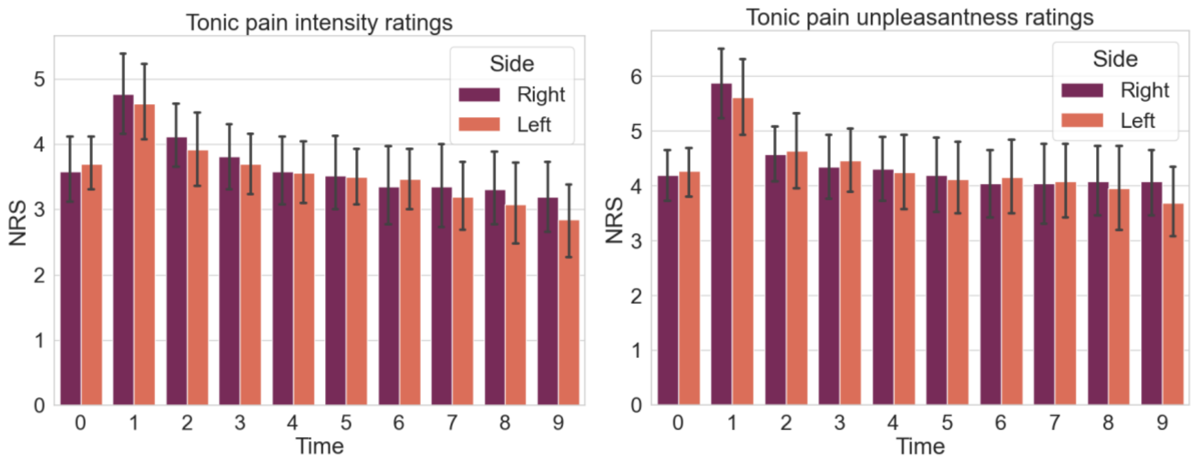


Figure 2. Plot of pressure pain ratings before the experiment (Time 0) and at 2-minute intervals throughout the extinction blocks. Ratings are averaged over each block.

**B -** Gaze Direction and Hand Position During Pain Anticipation

Gaze direction during each trial was extracted. Instances exceeding 2 SDs for 150 consecutive data points were removed and linearly interpolated. Trials containing >= 25% interpolated data were removed from further analysis (no trials exceeded this threshold). Interpolated data were subject to a median filter (kernel = 5), and the baseline period was removed. The number of fixations on the looming rock was extracted from the last 500 ms of the trial and averaged for each condition. Normalised pupil gaze direction over the entire trial duration showed a point of gaze origin directed slightly towards the right at the start of the trial, followed by a modest leftward shift after cue offset, which returned to the midpoint by the end of the trial. Results showed no statistically significant effects of cue, block or timebin on gaze direction ($p > .05$). Individual LMMs were used to predict changes in gaze direction with tonic pain, cue congruency and timebin. The main effects of cue congruency and timebin were not significant, nor were there any significant cue × timebin interactions (all $p > .05$; Supplementary Materials XX)

**Table 2. Fixed effects for gaze direction.**

| **Measure** | **Predictor** | **b** | **SE** | **df** | **t value** | ***p*** |
|---|---|---|---|---|---|---|
| **Gaze Direction** | Intercept | 0.66 | 2.46 | 29.46 | 0.27 | 0.791 |
| *(Gaze Direction ~* | Cue | -0.18 | 0.76 | 1184.97 | -0.23 | 0.819 |
| *Cue * Block ** | Block | 0.02 | 0.62 | 1185.00 | 0.03 | 0.975 |
| *Timebin)* | Timebin | -0.15 | 0.23 | 1184.97 | -0.66 | 0.511 |
| | Cue × Block | -0.11 | 0.76 | 1184.97 | -0.14 | 0.886 |
| | Cue × Timebin | 0.06 | 0.28 | 1184.97 | 0.21 | 0.831 |
| | Block × Timebin | -0.09 | 0.23 | 1184.97 | -0.42 | 0.677 |
| | Cue × Block × Timebin | 0.07 | 0.28 | 1184.97 | 0.25 | 0.805 |
| **Gaze Direction** | Intercept | 0.24 | 2.40 | 26.28 | 0.10 | 0.920 |
| *(Gaze Direction ~ TonicSide + Timebin)* | Congruency | 0.03 | 0.36 | 772.97 | 0.09 | 0.932 |

Hand position from left and right hands was extracted from trial data along *x, y* and *z* dimensions and normalised to the starting point for each block. Left- and right-hand positions were averaged to get an overall hand position value for each block, cue type and tonic pain side (for extinction data). Changes in hand position in *x, y* and *z* (forward-backwards; up-down; left-right) dimensions were assessed during each condition. Mean displacement during the experiment was *x* -8.5 ± 20.5 mms, *y* -6.04 ± 11.96 mms, *z* 2.56 ± 10.94 mms. A repeated measures ANOVA found no significant difference in mean displacement in any dimension throughout the experiment ($p > .05$).

**C – EEG data processing**

EEG data were processed using EEGLab [46]. Continuous EEG data were split into 8 s epochs (-2.5–5.5 s around cue onset). Data were re-referenced to the common average [47] and filtered from 1–70 Hz. The CleanLine plugin was used to estimate and remove sinusoidal line noise. Due to the presence of line noise in signal power spectra after executing CleanLine, an additional notch filter from 48–52 Hz was also implemented. Data for all 4 conditions were subsequently merged, resulting in one datafile per participant.

Oculomotor and muscular artefacts were removed from the data using independent component analysis (ICA). For optimal ICA [48], a subset of the data filtered from 1–30 Hz was created for each participant. RUNICA was conducted on the streamlined datafile, and ICA weights were exported and back-projected onto the original merged datafile. Data were epoched from -2.5–3.5 s around cue onset. Electrode channels with large artefacts were identified with visual inspection and interpolated to a maximum of 10% of all electrodes. Epochs containing extreme data due to movement or muscle artefacts were excluded using a semi-automated method. Outlier values were identified using *pop_eegthresh.m* with limits of -125–125 µV during the entire epoch. Improbable data were marked using *pop_jointprob.m* using single channel and global channel limits of 7 SDs. All data were visually inspected and marked trials manually reviewed. The average number of epochs remaining after artefact correction for each block were: conditioning 69.56 ± 3.55, extinction 104.06 ± 3.68. Accepted trials were not significantly different between cue types ($F(2,309) = .013$, $p = .987$).

To enhance small local currents and arrive at reference-free data, data were transformed into current source density using a Laplacian Spherical Spline interpolation method [49,50] implemented in FieldTrip [51] (http://fieldtriptoolbox.org) using default regularisation (lambda 1e-5) and skin conductivity (0.33 S/m) parameters and 9 degrees of Legendre polynomials, as recommended by FieldTrip for datasets with <= 32 channels. Power spectra were computed using a discrete Fourier time-frequency transformation. Power spectral densities were computed using Welch's method in 1 s windows shifted in overlapping 0.1 s segments to yield a power time series of 61 points. Data were smoothed using a 4 Hz multitaper. Spectral power was estimated from 1–70 Hz with a frequency resolution of 1 Hz.

**D** – Covariate analyses

The potential spatial hemisphere bias for the tonic pain-cue congruency effect was investigated in EEG data. For assessing the role of electrode hemisphere (contralateral and ipsilateral to the cue) on the pattern of EEG band power, an additional hemisphere factor was included for symmetrical electrodes. As smaller clusters are likely to have smaller degrees of freedom, a broader frontal cluster was used for the assessment of hemisphere lateralisation encompassing F7, F3, F4 and

F8, and the occipital region was not included. No lateralised ERD/S effect of cue congruency were observed following FWE-correction (see Supplementary Materials Table 3).

To examine the additive explanatory value of physiological data to EEG band power changes, pupil diameter and gaze direction were added to LMMs with ERD. Gaze direction did not significantly predict the changes in EEG power spectra during extinction blocks following pain-related cues in any frequency band. Pupil diameter significantly predicted parietal ($F(2,25) = -2.9$, $p < .001$) and occipital beta ($F(2,25) = 19.08$, $p < .001$) and parietal theta ($F(2,25) = 19.08$, $p < .001$) band power. Following FWE-correction, no interactions were found between cue congruency and pupil diameter.

**Table 3. EEG fixed effects for Tonic pain – cue congruency with covariates**

| **Band** | **Predictor** | **Covariate** | **Cluster** | **b** | **SE** | **df** | **t value** | **p** |
|---|---|---|---|---|---|---|---|---|
| **Alpha** *(EEG ~ TonicPain * Covariate)* | Congruency | Intensity | Frontal | -8.68 | 4.10 | 884.00 | -2.12 | 0.034 |
| | Congruency × Intensity | Intensity | Frontal | 2.51 | 1.08 | 884.00 | 2.31 | 0.021 |
| | Intensity | Intensity | Frontal | -5.68 | 1.82 | 54.13 | -3.12 | 0.003* |
| | | | Parietal | -5.91 | 2.09 | 45.02 | -2.83 | 0.007* |
| | Congruency | Unpleasantness | Frontal | -13.50 | 4.13 | 884.00 | -3.27 | 0.001* |
| | | | Central | -6.32 | 2.43 | 2132.00 | -2.60 | 0.009* |
| | Congruency × Unpleasantness | Unpleasantness | Frontal | 3.19 | 0.91 | 884.00 | 3.52 | 0.000* |
| | | | Central | 1.56 | 0.54 | 2132.00 | 2.91 | 0.004* |
| | Unpleasantness | Unpleasantness | Frontal | -5.33 | 1.54 | 48.36 | -3.46 | 0.001* |
| | | | Central | -3.28 | 1.28 | 36.89 | -2.57 | 0.014* |
| | Congruency × Pupil Diameter | Pupil Diameter | Parietal | 21.27 | 10.33 | 2787.48 | 2.06 | 0.040 |
| | Pupil Diameter | Pupil Diameter | Frontal | -46.18 | 9.51 | 927.51 | -4.86 | 0.000* |
| | Congruency × Unpleasantness × Hemisphere | Unpleasantness + Hemisphere | Frontal | 2.58 | 1.30 | 1820 | 1.99 | 0.047 |
| | Congruency × Hemisphere | Unpleasantness + Hemisphere | Frontal | -11.61 | 5.90 | 1820 | -1.97 | 0.049 |
| **Beta** *(EEG ~ TonicPain * Covariate)* | Congruency | Unpleasantness | Parietal | -6.64 | 1.52 | 2756.00 | -4.36 | 0.000* |
| | | | Occipital | -6.36 | 2.14 | 884.00 | -2.97 | 0.003* |
| | | Unpleasantness + Hemisphere | Parietal | -6.76 | 2.26 | 2444 | -2.99 | 0.003 |
| | | | Central | -5.42 | 2.48 | 1820 | -2.19 | 0.029 |
| | Congruency × Hemisphere | Pupil Diameter + Hemisphere | Central | -3.37 | 1.19 | 1819.85 | -2.84 | 0.005 |
| | | | Occipital | 4.49 | 1.84 | 571.95 | 2.44 | 0.015 |
| | | Unpleasantness | Parietal | 1.42 | 0.34 | 2756.00 | 4.24 | 0.000* |

| | | | | | | | | |
|---|---|---|---|---|---|---|---|---|
| | Congruency × Unpleasantness | | Occipital | 1.27 | 0.47 | 884.00 | 2.69 | 0.007* |
| | Unpleasantness | Unpleasantness | Central | -1.22 | 0.56 | 64.98 | -2.16 | 0.034 |
| | | | Parietal | -1.81 | 0.59 | 47.85 | -3.06 | 0.004* |
| | | | Occipital | -1.47 | 0.71 | 49.57 | -2.07 | 0.044 |
| | Unpleasantness × Hemisphere | Unpleasantness + Hemisphere | Central | 1.12 | 0.54 | 1820 | 2.05 | 0.040 |
| | Congruency × Pupil Diameter | Pupil Diameter | Central | -11.19 | 5.32 | 2171.19 | -2.10 | 0.036 |
| | Pupil Diameter | Pupil Diameter | Central | 8.33 | 3.89 | 2157.72 | 2.14 | 0.032 |
| | | | Parietal | -11.30 | 3.54 | 2508.66 | -3.20 | 0.001 |
| | | | Occipital | -17.17 | 4.90 | 919.23 | -3.50 | 0.000* |
| | Congruency × Unpleasantness | Unpleasantness + Hemisphere | Central | 1.55 | 0.54 | 1820 | 2.84 | 0.005 |
| | | | Parietal | 1.19 | 0.50 | 2444 | 2.40 | 0.016 |
| | Congruency × Unpleasantness × Hemisphere | Unpleasantness + Hemisphere | Central | -1.92 | 0.77 | 1820 | -2.49 | 0.013 |
| | Congruency × Pupil Diameter × Hemisphere | Pupil Diameter + Hemisphere | Central | -24.73 | 10.40 | 1820 | -2.38 | 0.018 |
| **Theta** *(EEG ~ TonicPain + Covariate)* | Pupil Diameter | Pupil Diameter | Frontal | -57.21 | 15.18 | 733.19 | -3.77 | 0.000* |

Note. Asterisks denote effects that survived FWE correction.

### D – Computational modelling of trial-by-trial anticipatory changes

#### Table 4. Fixed effects for pupil dilation – Pavlovian learning

Reference level for congruency: incongruent.

| Measure | Predictor | Timebin | b | SE | t value | p |
|---|---|---|---|---|---|---|
| **Pupil Diameter** *(Pupil Diameter ~ V * Timebin)* | Intercept | | -0.01 | 0.003 | -2.303 | 0.021 |
| | Timebin | Early | -0.1 | 0.005 | -19.323 | 0.000* |
| | | Late | -0.04 | 0.005 | -8.181 | 0.000* |
| | | Rock | 0 | 0.005 | -0.288 | 0.773 |
| | V | | 0 | 0.009 | -0.059 | 0.953 |
| | V × Timebin | Early | 0.02 | 0.013 | 1.513 | 0.13 |
| | | Late | 0.06 | 0.013 | 4.862 | 0.000* |
| | | Rock | 0.1 | 0.013 | 8.099 | 0.000* |
| **Pupil Diameter** *(Pupil Diameter ~ V*Congruency * Timebin)* | Intercept | | -0.02 | 0.008 | -2.187 | 0.029 |
| | Timebin | Early | -0.1 | 0.011 | -9.424 | 0.000* |
| | | Late | -0.04 | 0.011 | -3.38 | 0.001* |
| | | Rock | 0 | 0.011 | 0.302 | 0.763 |
| | Congruency | | 0.01 | 0.011 | 0.717 | 0.473 |
| | V | | 0.01 | 0.033 | 0.346 | 0.729 |
| | V × Congruency | | 0 | 0.047 | 0.091 | 0.927 |
| | V × Timebin | Early | 0.08 | 0.046 | 1.821 | 0.069 |
| | | Late | 0.03 | 0.046 | 0.627 | 0.53 |
| | | Rock | 0.07 | 0.046 | 1.413 | 0.158 |
| | V × Timebin × Congruency | Early | 0.01 | 0.015 | 0.466 | 0.641 |
| | | Late | 0 | 0.015 | -0.087 | 0.931 |
| | | Rock | 0 | 0.015 | -0.05 | 0.96 |

#### Table 5. Fixed effects for EEG – Pavlovian learning

Formula: EEG ~ V * Cluster. Reference level for congruency: incongruent.

| Predictor | Band | Timebin | Location | b | SE | t value | p |
|---|---|---|---|---|---|---|---|
| **Intercept** | | | | - 1.75 | 0.120 | -14.628 | 0.000* |
| **V** | | | | 0.14 | 0.288 | 0.479 | 0.632 |
| **Cluster** | Theta | Cue | Frontal | 10.04 | 0.495 | 20.266 | 0.000* |
| | | | Central | 11.05 | 0.494 | 22.380 | 0.000* |
| | | Early | Frontal | 3.40 | 0.496 | 6.846 | 0.000* |
| | | | Central | 1.44 | 0.493 | 2.915 | 0.004* |
| | | Late | Frontal | -4.80 | 0.496 | -9.687 | 0.000* |
| | | | Central | 1.04 | 0.493 | 2.106 | 0.035 |
| | Alpha | Cue | Central | 1.19 | 0.494 | 2.420 | 0.016* |

| | | | | | | | |
|---|---|---|---|---|---|---|---|
| | | | Parietal | -4.91 | 0.495 | -9.928 | 0.000* |
| | | Early | Central | -1.72 | 0.494 | -3.490 | 0.000* |
| | | | Parietal | -7.63 | 0.495 | -15.416 | 0.000* |
| | | Late | Central | 0.88 | 0.495 | 1.788 | 0.074 |
| | | | Parietal | -1.68 | 0.498 | -3.382 | 0.001* |
| | Beta | Cue | Central | 0.62 | 0.492 | 1.260 | 0.208 |
| | | | Parietal | -5.61 | 0.492 | -11.398 | 0.000* |
| | | Early | Central | -0.35 | 0.492 | -0.713 | 0.476 |
| | | | Parietal | -0.99 | 0.492 | -2.007 | 0.045 |
| | | Late | Central | -0.85 | 0.492 | -1.727 | 0.084 |
| **V × Cluster** | Theta | Cue | Frontal | 5.40 | 1.193 | 4.526 | 0.000* |
| | | | Central | 6.14 | 1.186 | 5.174 | 0.000* |
| | | Early | Frontal | 5.23 | 1.194 | 4.384 | 0.000* |
| | | | Central | 2.53 | 1.186 | 2.130 | 0.033 |
| | | Late | Frontal | 1.07 | 1.190 | 0.895 | 0.371 |
| | | | Central | -0.74 | 1.187 | -0.625 | 0.532 |
| | Alpha | Cue | Central | 3.19 | 1.188 | 2.687 | 0.007* |
| | | | Parietal | 6.00 | 1.190 | 5.045 | 0.000* |
| | | Early | Central | -2.92 | 1.187 | -2.458 | 0.014* |
| | | | Parietal | -1.73 | 1.189 | -1.457 | 0.145 |
| | | Late | Central | -6.33 | 1.188 | -5.324 | 0.000* |
| | | | Parietal | -5.07 | 1.195 | -4.239 | 0.000* |
| | Beta | Cue | Central | 0.27 | 1.184 | 0.225 | 0.822 |
| | | | Parietal | 0.90 | 1.184 | 0.763 | 0.445 |
| | | Early | Central | -2.76 | 1.184 | -2.333 | 0.020* |
| | | | Parietal | -1.73 | 1.184 | -1.458 | 0.145 |
| | | Late | Central | -4.89 | 1.185 | -4.127 | 0.000* |

### Table 6. Fixed effects for EEG – Pavlovian learning, Congruency Effects

Formula: EEG ~ V * Congruency * Cluster. Reference level for congruency: incongruent.

| **Predictor** | **Band** | **Timebin** | **Location** | **b** | **SE** | **t value** | **p** |
|---|---|---|---|---|---|---|---|
| **Intercept** | | | | -2.48 | 0.253 | -9.824 | 0.000* |
| **Congruency** | | | | -0.47 | 0.357 | -1.328 | 0.184 |
| **V** | | | | -2.26 | 1.251 | -1.806 | 0.071 |
| **V × Congruency** | | | | 5.58 | 1.795 | 3.111 | 0.002* |
| **Cluster** | Theta | Cue | Frontal | 10.74 | 1.047 | 10.262 | 0.000* |
| | | | Central | 10.74 | 1.043 | 10.294 | 0.000* |
| | | Early | Frontal | 2.66 | 1.049 | 2.537 | 0.011 |
| | | | Central | 0.85 | 1.04 | 0.816 | 0.414 |
| | | Late | Frontal | -5.89 | 1.049 | -5.617 | 0.000* |
| | | | Central | 0.28 | 1.042 | 0.269 | 0.788 |
| | Alpha | Cue | Central | 1.39 | 1.041 | 1.338 | 0.181 |

| | | | | | | | |
|---|---|---|---|---|---|---|---|
| | | | Parietal | -3.76 | 1.043 | -3.605 | 0.000* |
| | | Early | Central | -2.07 | 1.039 | -1.99 | 0.047 |
| | | | Parietal | -6.69 | 1.044 | -6.41 | 0.000* |
| | | Late | Central | 0.93 | 1.043 | 0.891 | 0.373 |
| | | | Parietal | -0.85 | 1.052 | -0.808 | 0.419 |
| | Beta | Cue | Central | -0.05 | 1.04 | -0.052 | 0.958 |
| | | | Parietal | -5.65 | 1.038 | -5.446 | 0.000* |
| | | Early | Central | -0.7 | 1.037 | -0.677 | 0.499 |
| | | | Parietal | -1.15 | 1.037 | -1.112 | 0.266 |
| | | Late | Central | -1.04 | 1.037 | -1.002 | 0.317 |
| **V × Cluster** | Theta | Cue | Frontal | 0.98 | 5.165 | 0.19 | 0.849 |
| | | | Central | 5.44 | 5.159 | 1.055 | 0.291 |
| | | Early | Frontal | 19.6 | 5.176 | 3.787 | 0.000* |
| | | | Central | 7.35 | 5.15 | 1.427 | 0.154 |
| | | Late | Frontal | 16.27 | 5.169 | 3.147 | 0.002* |
| | | | Central | 3.41 | 5.162 | 0.66 | 0.509 |
| | Alpha | Cue | Central | -2.34 | 5.157 | -0.454 | 0.65 |
| | | | Parietal | -3.47 | 5.16 | -0.673 | 0.501 |
| | | Early | Central | -6.15 | 5.149 | -1.195 | 0.232 |
| | | | Parietal | -9.19 | 5.153 | -1.784 | 0.074 |
| | | Late | Central | -18.3 | 5.18 | -3.535 | 0.000* |
| | | | Parietal | -16.15 | 5.182 | -3.117 | 0.002* |
| | Beta | Cue | Central | 3.49 | 5.153 | 0.678 | 0.498 |
| | | | Parietal | 10.72 | 5.149 | 2.083 | 0.037 |
| | | Early | Central | -7.06 | 5.151 | -1.371 | 0.171 |
| | | | Parietal | 1.01 | 5.151 | 0.197 | 0.844 |
| | | Late | Central | -2.5 | 5.149 | -0.485 | 0.628 |
| **Cluster × Congruency** | Theta | Cue | Frontal | -4.57 | 1.476 | -3.095 | 0.002* |
| | | | Central | -0.29 | 1.47 | -0.2 | 0.841 |
| | | Early | Frontal | -1.76 | 1.479 | -1.192 | 0.233 |
| | | | Central | 0.43 | 1.468 | 0.292 | 0.77 |
| | | Late | Frontal | -0.81 | 1.479 | -0.547 | 0.585 |
| | | | Central | 1.83 | 1.47 | 1.244 | 0.214 |
| | Alpha | Cue | Central | 0.91 | 1.469 | 0.616 | 0.538 |
| | | | Parietal | -1.66 | 1.473 | -1.128 | 0.259 |
| | | Early | Central | 0.94 | 1.468 | 0.639 | 0.523 |
| | | | Parietal | -0.07 | 1.474 | -0.046 | 0.963 |
| | | Late | Central | 0.8 | 1.472 | 0.547 | 0.585 |
| | | | Parietal | 0.09 | 1.483 | 0.061 | 0.951 |
| | Beta | Cue | Central | 1.58 | 1.466 | 1.077 | 0.281 |
| | | | Parietal | 0.43 | 1.465 | 0.293 | 0.769 |
| | | Early | Central | 1.61 | 1.464 | 1.102 | 0.27 |

| | | | | | | | |
|---|---|---|---|---|---|---|---|
| | | | Parietal | 0.75 | 1.463 | 0.513 | 0.608 |
| | | Late | Central | 0.86 | 1.465 | 0.588 | 0.557 |
| **V × Cluster × Congruency** | Theta | Cue | Frontal | 16.7 | 7.405 | 2.255 | 0.024 |
| | | | Central | -2.36 | 7.394 | -0.319 | 0.75 |
| | | Early | Frontal | 5.67 | 7.436 | 0.762 | 0.446 |
| | | | Central | -3.4 | 7.396 | -0.459 | 0.646 |
| | | Late | Frontal | -1.49 | 7.444 | -0.2 | 0.841 |
| | | | Central | -12.28 | 7.402 | -1.659 | 0.097 |
| | Alpha | Cue | Central | -0.78 | 7.398 | -0.105 | 0.916 |
| | | | Central | 8.07 | 7.406 | 1.089 | 0.276 |
| | | Early | Parietal | -0.22 | 7.392 | -0.03 | 0.976 |
| | | | Central | 0.24 | 7.397 | 0.032 | 0.974 |
| | | Late | Parietal | 4.45 | 7.413 | 0.6 | 0.548 |
| | | | Central | -3.41 | 7.425 | -0.459 | 0.646 |
| | Beta | Cue | Central | -2.8 | 7.395 | -0.379 | 0.705 |
| | | | Parietal | -1.78 | 7.388 | -0.24 | 0.81 |
| | | Early | Central | 3.15 | 7.4 | 0.426 | 0.67 |
| | | | Parietal | -1.39 | 7.388 | -0.188 | 0.851 |
| | | Late | Central | -4.3 | 7.386 | -0.583 | 0.56 |
